# A novel function for endothelial protease-activated receptors in modulating insulin receptor activity with implications for diabetes

**DOI:** 10.1101/2025.03.21.644607

**Authors:** Rahul Rajala, Courtney T. Griffin

## Abstract

Thrombin, a serine protease with increased activity in diabetics, signals through protease-activated receptors 1 and 4 (PAR1/PAR4) on endothelial cells (ECs). While studying the roles of endothelial PAR1/4 in diabetic pathology, we found that mice with inducible deletion of both receptors on ECs (*Par1/4^iECko^*) displayed increased insulin sensitivity and were protected against streptozotocin (STZ)-induced diabetes. Concordantly, we found that cultured primary ECs with PAR1/4 deficiency had increased basal activity/phosphorylation of the insulin receptor (IR) and insulin transcytosis. This elevated IR activity correlated with reduced activity of protein tyrosine phosphatase 1B (PTP1B), which is a negative regulator of IR activity. Lastly, *Par1/4^iECko^*mice with additional deletion of one allele of the IR gene demonstrated restoration of diabetic phenotypes after STZ treatment, indicating that these phenotypes are driven by heightened IR activity. These findings establish a novel link between endothelial PAR signaling and IR regulation, underscoring the critical role of ECs in metabolic homeostasis and identifying a potential therapeutic target for diabetes.

**GRAPHICAL ABSTRACT:** 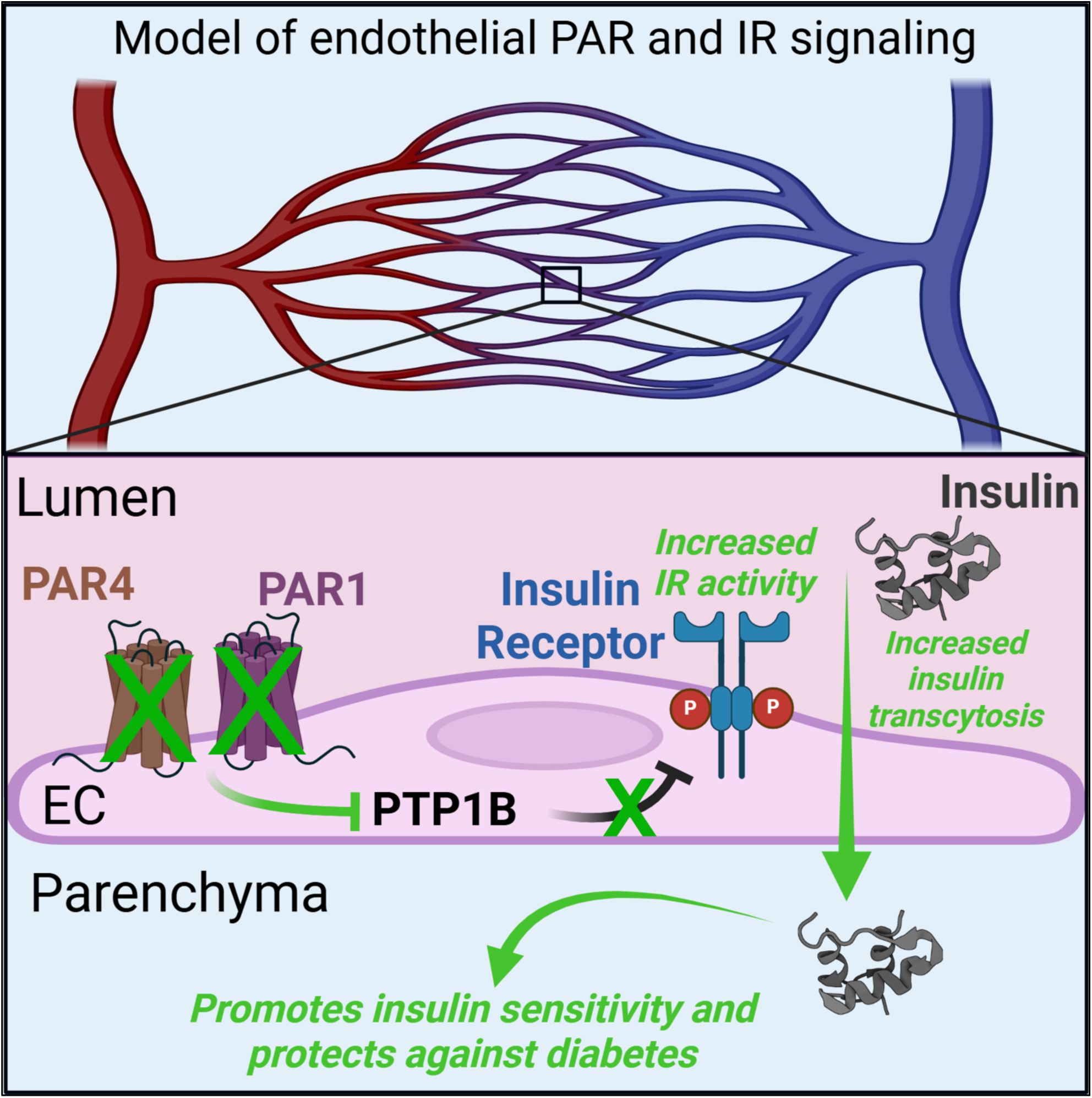

## INTRODUCTION

The vascular endothelium which lines the lumen of blood vessels, is an expansive layer of cells that covers a total surface area of 270-720 square meters (1–3). The endothelium is a key component of the vasculature with multiple functions, including trafficking hormones and nutrients to target tissues and regulating hemostasis, inflammation, and vascular permeability (2–4). With such crucial and varied functions, it is unsurprising that endothelial dysfunction worsens different pathological conditions in specific organs.

Thrombin, a serine protease, is involved in the development and maintenance of the vasculature (5). Thrombin signals through protease-activated receptors 1 and 4 (PAR1/PAR4), which are G protein-coupled receptors (GPCRs) found on endothelial cells (ECs) as well other cell types (2, 3). Thrombin has a different affinity for PAR1 (50 pM) and PAR4 (5000 pM), as well as different kinetics for signaling and desensitization (3). PAR signaling in ECs is implicated in a variety of physiological processes, and its dysregulation is linked to pathology (2, 3, 5, 6). Recent studies from our lab demonstrated pathological roles for endothelial PAR1/4 in an acute model of thrombin generation. However, the contribution of these receptors to diseases with more moderate but chronic levels of thrombin is unknown.

Diabetes is a prevalent disease characterized by hyperglycemia. Although the disease is predominantly described as a metabolic disorder, it also presents with numerous cardiovascular complications including gradual increases in thrombin generation leading to hypercoagulation (7). Thrombin inhibitors have beneficial effects in diabetic mouse models, including reduced expression of inflammatory cytokines and adhesion molecules in the vasculature (8). Independently, *Par1* and *Par4* expression have been shown to increase in the vasculature of diabetic mice (8); however, the cell-specific roles of PAR1 and PAR4 in blood vessels during diabetes remain a mystery. Our current study uses EC-specific PAR knockout mice to examine how endothelial PAR1 and PAR4 impact diabetic pathology.

## RESULTS

### Mice with deletion of endothelial *Par1/4* display significantly reduced hyperglycemia and other diabetic phenotypes after streptozotocin (STZ) challenge

To determine how diabetic pathology was impacted by the loss of endothelial PAR1 and PAR4 we generated mice with deletions of endothelial *Par1*, *Par4*, and *Par1/Par4*, respectively (hereafter, *Par1^iECko^*, *Par4^iECko^*, and *Par1/4^iECko^*) using *Par* floxed alleles crossed to the tamoxifen-inducible *Cdh5(PAC)-Cre^ERT2^* line, as described previously (2). This approach ensured selective deletion of the receptors in ECs of adult mice and avoided the lethality that occurs when endothelial *Par1/4* are deleted during embryonic development (5). We previously validated that this approach selectively removes *Par* expression in ECs in adult mice while preserving *Par* expression in parenchymal tissues (2). To generate diabetic *Par^iECko^* mice and suitable Cre-negative littermate controls, all experimental mice were treated with tamoxifen to induce gene deletion followed by STZ to induce hyperglycemia (blood glucose > 250 mg/dL) (**Figure 1A**). STZ is an alkylating agent that selectively ablates insulin-producing beta cells in the pancreatic islets of Langerhans, resulting in a mouse model of Type I diabetes with hyperglycemia and inflammation (9).

**Figure 1.**
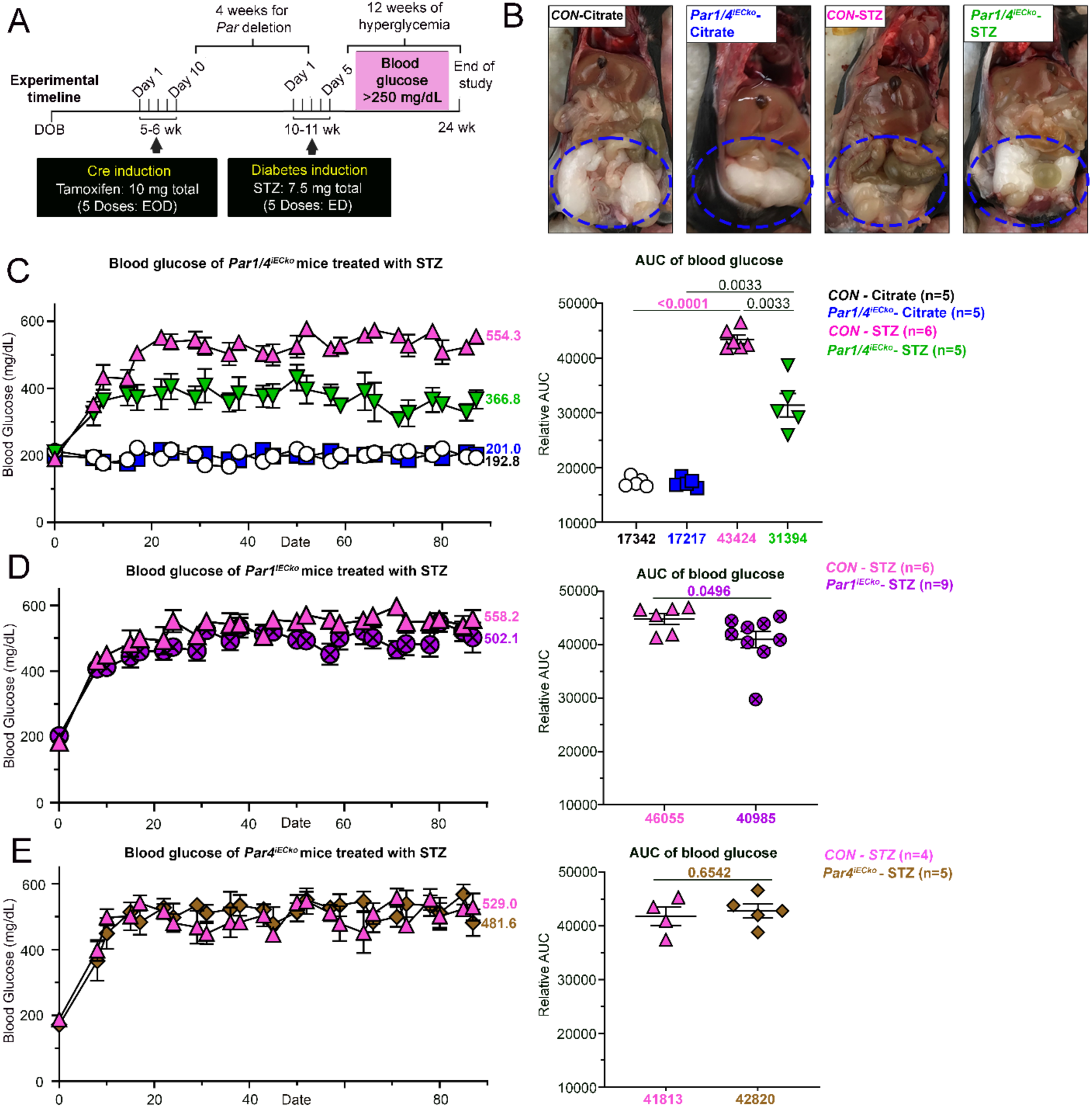
Dual loss of endothelial PAR1 and PAR4 protects mice against STZ-induced hyperglycemia. **(A)** Outline of experimental procedures performed on mice. DOB: date of birth; EOD: every other day; ED: every day. **(B)** Representative images of abdominal fat pads in control and *Par1/4^iECko^* mice at the end of the study. **(C)** Blood glucose levels and area under the curve (AUC) of STZ- and citrate-treated male *Par1/4^iECko^*, **(D)** *Par1^iECko^*, **(E)** and *Par4^iECko^* mice compared to littermate controls (CON). Mice were challenged with 7.5 mg of STZ, and diabetes was allowed to progress for 12 weeks. For (B, D, E), blood glucose levels are shown in mg/dL. Final measurements taken at the end of the study are displayed on the respective graphs. <u>Statistics</u>: (C) Analyzed by Welch ANOVA followed by a 2-stage linear step-up procedure of Benjamini, Krieger, and Yekutieli for multiple comparison testing; q values are displayed in the graphs. (D) Analyzed by Mann-Whitney U test followed by a 2-stage linear step-up procedure of Benjamini, Krieger, and Yekutieli for multiple comparison testing; p values are displayed in the graphs. (E) Analyzed by unpaired t-test followed by a 2-stage linear step-up procedure of Benjamini, Krieger, and Yekutieli for multiple comparison testing; p values are displayed in the graphs. All scatter dot plots show mean ± SEM.

We found that *Par1/4^iECko^* mice were protected against STZ-induced hyperglycemia (**Figure 1C**). STZ-treated *Par1/4^iECko^*mice displayed significantly reduced blood glucose levels (∼27.8%, p<0.0012) compared to control littermates. STZ-treated *Par1/4^iECko^*mice also had larger abdominal fat pads than littermate controls, which displayed diabetic lipoatrophy (**Figure 1B**). Concordantly, STZ-treated *Par1/4^iECko^* mice showed higher normalized body weights (∼7%) compared to littermate controls (**Figure S1**). This protection against STZ-induced hyperglycemia was not observed in single-knockout mice (**Figure 1D&E**). STZ-treated *Par1^iECko^* did display marginally reduced blood glucose levels (∼8.8%, p<0.0496) compared to control littermates (**Figure 1D**), but this effect was minimal when compared to *Par1/4^iECko^* mice (**Figure S2**).

Based on these initial findings, we assessed additional phenotypes associated with hyperglycemia. We assessed glucose metabolism indirectly by measuring hyaluronan (HA) levels. HA is an extracellular matrix glycosaminoglycan composed of repeating disaccharides of glucuronic acid and N-acetylglucosamine (**Figure S3A**). The synthesis of these constituent disaccharides is dependent on insulin-mediated cellular uptake and metabolism of glucose (10). In the context of severe and persistent hyperglycemia, we found significantly reduced serum HA levels in control mice (∼43%, p<0.0047) (**Figure S3B**). However, this reduction was attenuated in STZ-treated *Par1/4^iECko^*mice, suggesting increased cellular internalization and metabolism of glucose in these mice.

Hyperglycemia also results in polyuria-mediated dehydration which is associated with elevated hematocrit and hyperalbuminemia (11). These phenotypes were observed in STZ-treated control mice but were attenuated in STZ-treated *Par1/4^iECko^* littermate (**Figure S3C&D**). Studies on diabetic patients have shown increased mean eosinophil counts (12). We observed a moderate increase in eosinophils in STZ-treated control mice that were attenuated in STZ-treated *Par1/4^iECko^* littermates (**Figure S4G**).

STZ-induced induced hyperglycemia results in renal hypertrophy, as measured by kidney-to-body weight (KBW) ratios (13). STZ-treated control mice showed increased KBW, which was attenuated in STZ-treated *Par1/4^iECko^* mice (**Figure S5B**). However, renal function itself was unaffected by our diabetic challenge, as STZ-treated mice showed minimal changes in serum creatinine and blood urea nitrogen (BUN) from vehicle-treated littermates (**Figure S5C&D**). Lastly, when we measured thrombin levels, STZ-treated mice showed marginally elevated thrombin levels as measured by thrombin-antithrombin (TAT) complexes in plasma (**Figure S6**). However, these levels were not significantly changed between STZ-treated *Par1/4^iECko^*mice and littermate controls.

Overall, we observed that STZ-treated *Par1/4^iECko^* mice had reduced hyperglycemia, increased body weights, increased fat pads, increased glucose metabolism, reduced dehydration, and reduced renal hypertrophy compared to littermate controls. This suggests that *Par1/4^iECko^* mice are protected against STZ-induced diabetes and its associated sequelae.

Given that *Par1/4^iECko^* mice were protected against STZ-induced diabetes, we sought to determine if reducing endothelial PAR1 and PAR4 could reverse hyperglycemia in STZ-treated mice. We modified our experimental protocol to take advantage of the tamoxifen-inducible deletion of *Par1* and *Par4* in ECs after the onset of STZ-induced hyperglycemia (**Figure S7**). Before tamoxifen administration, the mice showed no significant difference in hyperglycemia (timepoint 1). Following tamoxifen induction, *Par1/4^iECko^* mice showed moderately reduced hyperglycemia from littermate controls (timepoint 2); *Par1/4^iECko^* mice also showed a significant reduction in blood glucose levels compared to their levels measured before gene deletion (timepoint 2 vs. 1; ∼18.7%, p<0.0478) (**Figure S7B&C**). However, these reductions in hyperglycemia in *Par1/4^iECko^* mice did not persist, and by the end of the study, they displayed hyperglycemia again (timepoint 3; **Figure S7**).

### *Par1/4^iECko^* mice display increased insulin sensitivity

Since STZ-induced diabetes is driven by a loss of pancreatic insulin production, we next sought to determine if increases in circulating insulin were responsible for *Par1/4^iECko^* mice being protected against STZ-induced hyperglycemia. We collected serum from mice at 12 weeks following the STZ challenge and measured circulating insulin levels by ELISA. We found ∼50% decreased levels of circulating insulin in STZ-treated control mice (**Figure 2A**), similar to previous studies (14). Circulating insulin levels were equivalently reduced in *Par1/4^iECko^* mice (**Figure 2A**). We next assessed insulin sensitivity in non-diabetic control and non-diabetic *Par1/4^iECko^* mice by performing insulin tolerance (**Figure 2B&C**) and glucose tolerance (**Figure 2D-E**) tests. *Par1/4^iECko^* mice showed significantly increased insulin sensitivity (14.3% reduced AUC, p<0.0149) and glucose tolerance (17.1% reduced AUC, p<0.0403) compared to littermate controls. These data suggest that *Par1/4^iECko^* mice can utilize insulin more efficiently.

**Figure 2.**
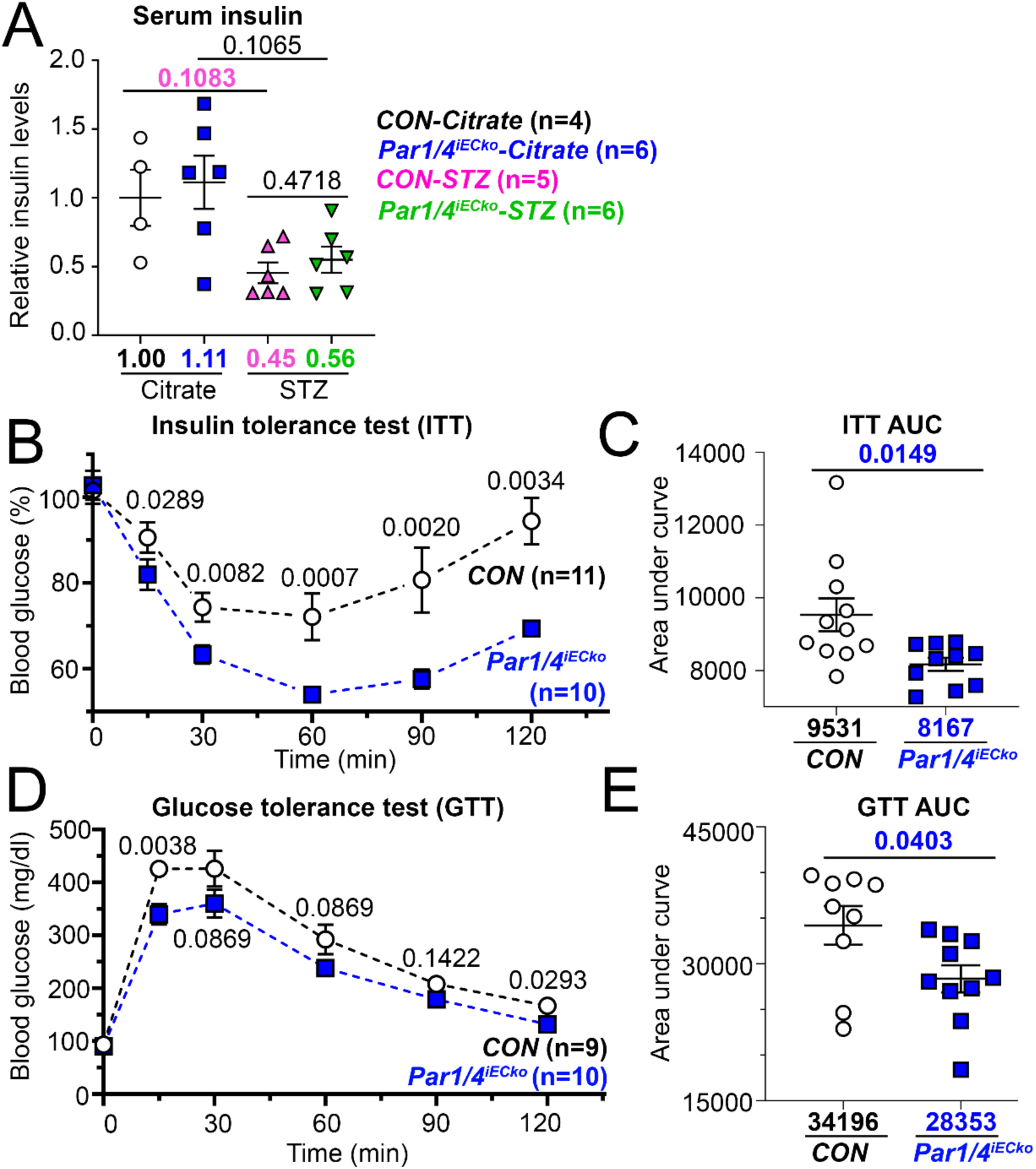
Dual loss of endothelial PAR1 and PAR4 increases insulin sensitivity in mice but not circulating insulin levels. **(A)** Serum insulin levels (ELISA) of mice which were collected after 6 hours of fasting. (n=4-6) **(B)** Insulin tolerance test (ITT) normalized to baseline reading (insulin administered: 1 U/kg). (**C**) The area under the curve (AUC) is calculated for each ITT measurement. (**D**) Glucose tolerance test (GTT) normalized to baseline reading (glucose administered: 1 g/kg). The AUC is calculated for each GTT measurement. <u>Statistics</u>: (A) Analyzed with a Welch ANOVA followed by a 2-stage linear step-up procedure of Benjamini, Krieger, and Yekutieli for multiple comparison testing; q values are displayed in the graphs. n=4–6 mice per group. (B, D) Analyzed with a Mann-Whitney U test followed by a 2-stage linear step-up procedure of Benjamini, Krieger, and Yekutieli for multiple comparison testing; q values are displayed in the graphs. (C, E) analyzed with a Welch t-test; p-values are displayed in the graphs. (B-E): n=9–11 mice per group. CON: control.

### PAR1/4-deficient ECs have increased basal insulin receptor activity and insulin transcytosis

Studies have shown that increased IR signaling in ECs promotes endothelial insulin transcytosis and insulin sensitivity (4, 15). Therefore, we sought to determine if loss of endothelial PAR1 and PAR4 increased IR activity and transcytosis in ECs, which could contribute to *Par1/4^iECko^* mice displaying improved whole body insulin sensitivity.

To determine if loss of endothelial PAR1 and PAR4 impacted endothelial IR signaling, we assayed primary Human Vein Umbilical Endothelial Cells (HUVECs), which do not express PAR4 (2). Using a siRNA for PAR1 that gave a (∼50-fold) decrease in *PAR1* expression (**Figure S8A)**, we generated PAR1/4-deficient ECs. We then used the uptake of a fluorescent glucose analog (2-NBDG) coupled with an IR inhibitor (HNMPA3) to assess IR signaling. We found that at baseline, PAR1/4-deficient ECs displayed 2-NBDG uptake levels approximately equivalent to levels observed in insulin-treated control ECs. Furthermore, this phenotype was attenuated with the addition of HNMPA3 (**Figure 3 and Figure S8B-D**). Similar attenuation of glucose uptake in PAR1/4-depleted ECs was also seen after treatment with a PI3K inhibitor (LY294002) (**Figure S8E**). Treatment of HUVECs with a PAR1 inhibitor (vorapaxar) (16) also resulted in an elevation of 2-NBDG uptake, although it was not as pronounced as with PAR1 knockdown (**Figure S8F**). These data suggest that loss of endothelial PARs increases basal IR activity.

**Figure 3.**
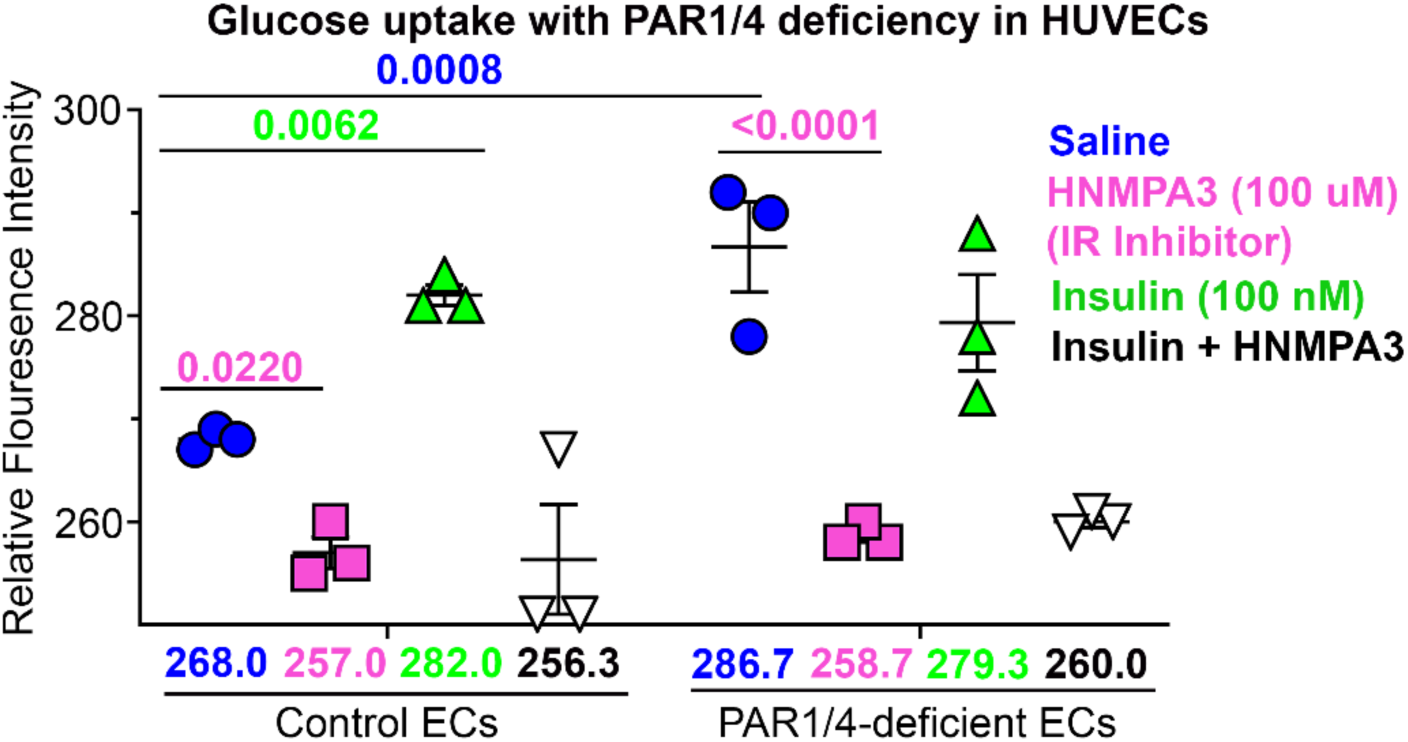
PAR1/4-deficient ECs have increased IR signaling. Flow cytometry analysis of uptake of 2-NBDG (a glucose analog) as a readout of IR signaling. Control and PAR1/4-deficient HUVECs were treated with saline (blue), an IR inhibitor (HNMPA3; pink), insulin (green), and insulin plus HNMPA (black). Analyzed by 3-way ANOVA followed by a 2-stage linear step-up procedure of Benjamini, Krieger, and Yekutieli for multiple comparison testing; q values are displayed in the graphs. n=3 technical replicates per group.

We next sought to determine whether PAR1/4-deficient ECs with heightened IR activity also had an increased ability to transcytose insulin. We plated a confluent monolayer of control and PAR1/4-deficient ECs on a transwell and measured the amount of biotinylated insulin that was transcytosed across the transwell by ELISA (**Figure 4A**). Treatment with an IR-blocking peptide (S931) reduced transcytosis of insulin, validating our assay (**Figure S9C**). In this experiment, PAR1/4-deficient ECs were generated using a lentiviral shRNA for PAR1 that yielded stable knockdown of *PAR1* expression (∼3.57 fold) (**Figure S9A&B**). A stable knockdown of PAR1 was employed since this experiment requires several days to ensure confluent monolayers form on the transwell. We found that PAR1/4-deficient ECs had significantly increased insulin transcytosis (p<0.0043) (**Figure 4B**). To investigate whether PAR1 activation conversely reduced transcytosis, we measured transcytosis following thrombin treatment but saw no difference compared to saline-treated controls (**Figure S9E**). Overall, these results suggest that loss of PAR1/4 increases basal IR activity and insulin transcytosis in ECs.

**Figure 4.**
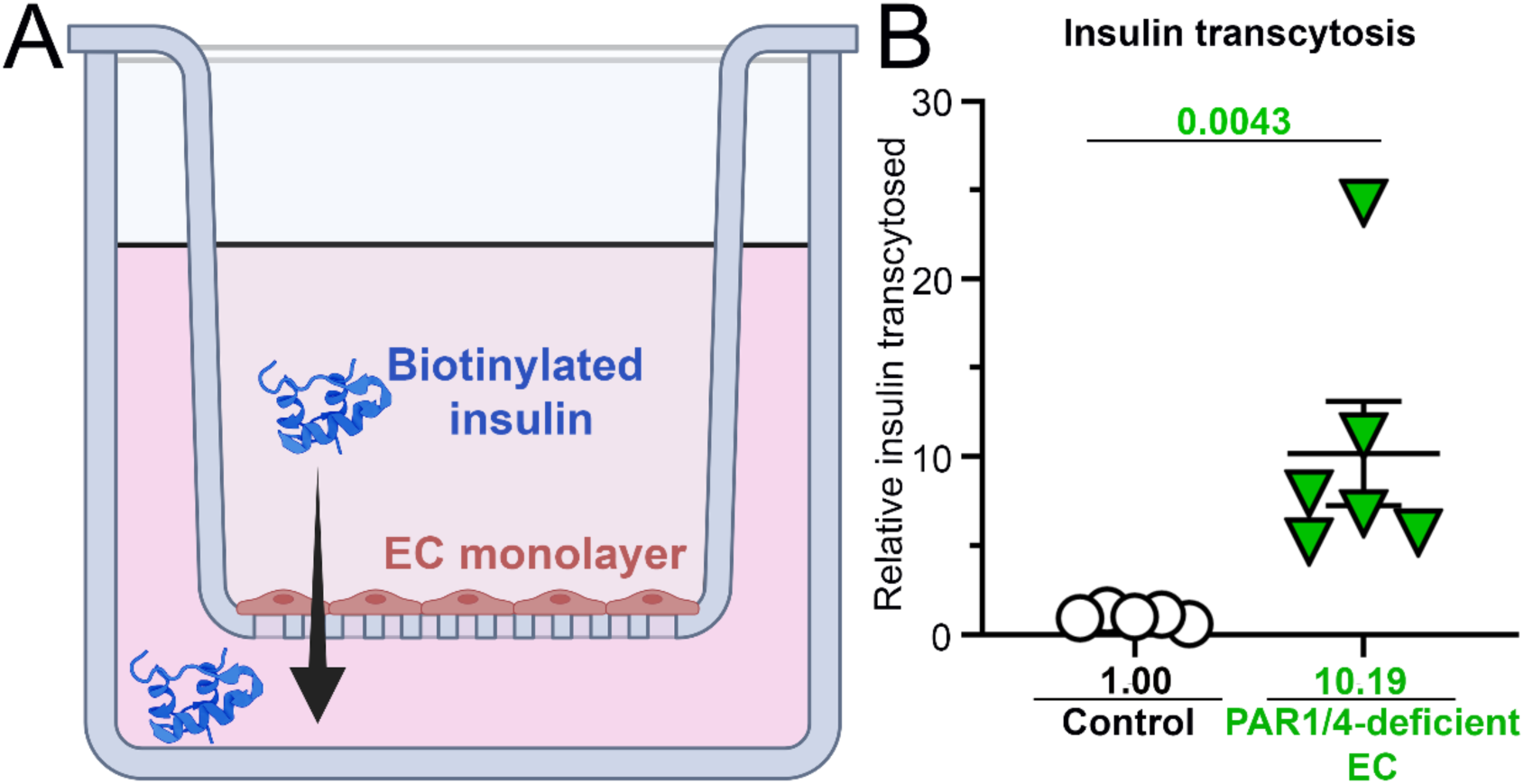
PAR1/4-deficient ECs have an increased ability to transcytose insulin. (**A**) Graphic of insulin transcytosis assay. (**B**) Relative levels of transcytosed insulin between control and PAR1/4-deficient HUVECs. Analyzed by Mann Whitney U-test; p values are displayed in the graphs. n=5-6 technical replicates per group.

### Increased insulin sensitivity in *Par1/4^iECko^* mice is unlikely to be related to endothelial IR splicing

Since PAR1/4*-*depleted EC displayed increased basal IR activity and insulin transcytosis, we sought to understand what drove these phenotypes. One possibility was a change in IR splicing. There are two spliceforms of IR: Type A (IR-A) and Type B (IR-B). These spliceforms are distinguished by the presence of a 12 amino acid fragment encoding exon 11, which is found in IR-B (**Figure S10A-D**) but is absent in IR-A (17). IR-A has higher basal activity (18), a higher binding affinity for insulin (**Figure S10E**) (19), and faster internalization kinetics (20) than IR-B (19). In PAR1/4-deficient ECs, we found significant increases in *INSR-A* and *INSR* transcript levels compared to control cells (**Figure S10F-G**), which could account for their heightened IR activity. However, when we isolated EC-specific mRNA using translating ribosome affinity purification (TRAP) from murine quadriceps, we found the opposite effect: loss of endothelial PAR1/4 in ECs *in vivo* significantly reduced expression of *Insr-A* (**Figure S11**). Although these combined *in vitro* and *in vivo* results suggest a relationship between endothelial PARs and IR splicing, the diametrically opposed spliceform expression patterns in these experiments indicate that changes in IR splicing are unlikely to drive increased insulin sensitivity in *Par1/4^iECko^* mice.

### PAR1/4-deficient ECs have increased basal IR phosphorylation

We next sought to determine if the increased basal IR activity in PAR1/4-deficient HUVECs was due to increased IR phosphorylation. Like all receptor tyrosine kinases, IR activation relies on trans-autophosphorylation of tyrosine residues on its c-terminal loops (21). p85 is the regulatory subunit of phosphoinositide 3-kinase (PI3K), which binds phosphotyrosine residues on RTKs via its Src Homology 2 (SH2) domains, so we performed p85-glutathione S-transferase (GST) pulldowns (22) in PAR1/4*-* deficient ECs to assess IR phosphorylation (**Figure 5A**). Using this approach, we isolated phosphorylated RTKs from ECs and immunoblotted for IR (**Figure S12**). This approach is preferred over the use of phosphospecific IR antibodies, which have strong cross-reactivity to phospho-IGF-1R. It is also preferred over traditional immunoprecipitation, which requires significant protein input.

**Figure 5.**
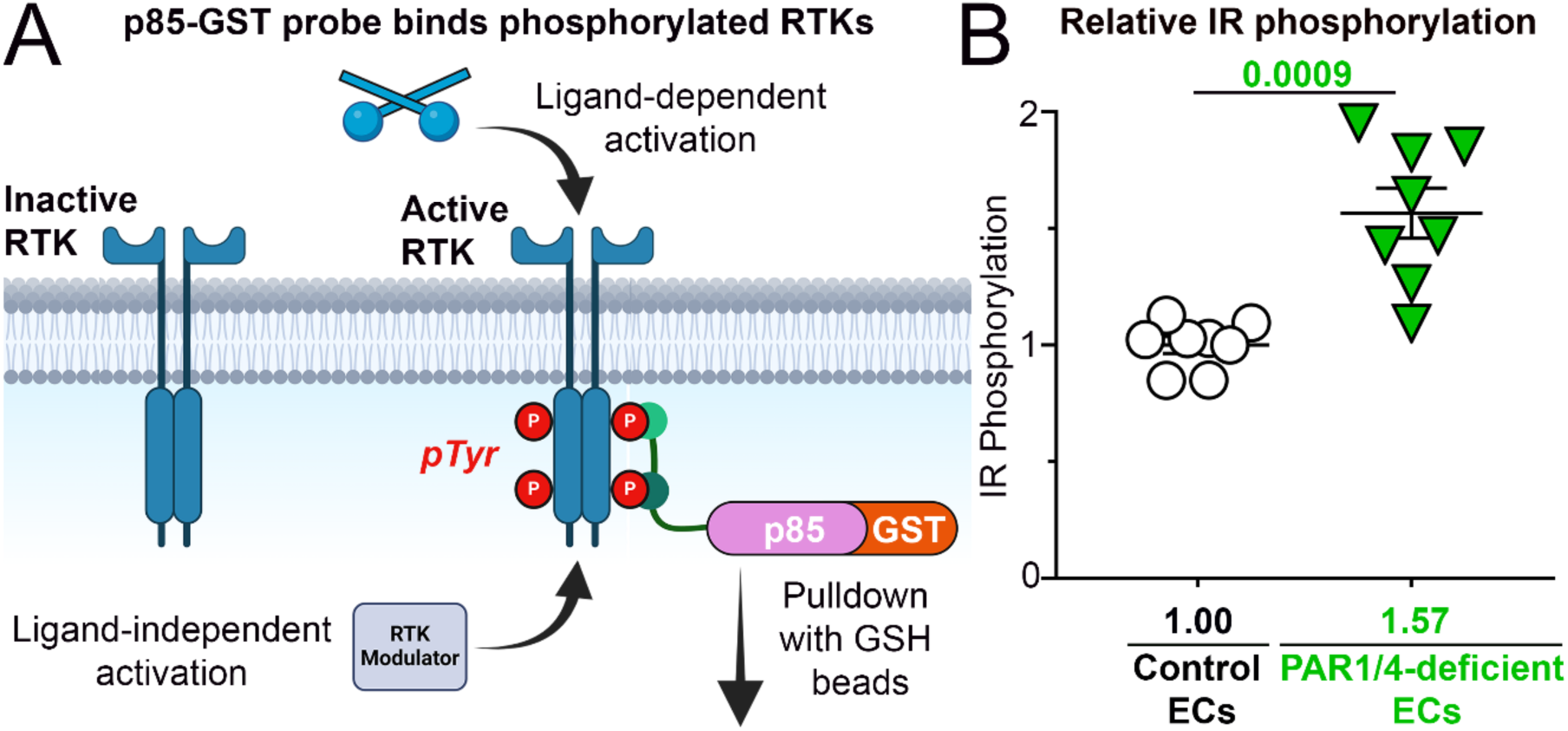
PAR1/4-deficient ECs have increased IR phosphorylation. (**A**) Schematic of p85-GST pulldown to measure IR phosphorylation. (pTyr – phosphotyrosine) (**B**) Quantification of relative IR phosphorylation in control and PAR1/4-deficient ECs. Analyzed by Welch’s t-test; p values are displayed in the graphs. n=8 replicates per group.

We found PAR1/4-deficient ECs had significantly increased basal IR phosphorylation (∼57%, p<0.0009) (**Figure 5B & Figure S13C**), with levels similar to ECs treated with insulin (∼105%, p<0.0002) (**Figure S13A&B**). ECs treated with an inhibitor of PAR1 (vorapaxar) showed no significant change in IR phosphorylation (p<0.5462) (**Figure S13A&D**). This is in line with our previous findings showing only marginal changes in IR activity with vorapaxar treatment. These data indicate that the loss (but not inhibition) of PARs drives IR phosphorylation. When we assayed total IR levels in PAR1/4-deficient ECs, we found total IR levels reduced by immunoblots and immunocytochemistry compared to control cells (**Figure S14**). This may be a result of receptor downregulation due to the heightened IR activity. Meanwhile, transcripts for the IR gene (*INSR*) were increased in PAR1/4-deficient ECs (**Figure S11G**), perhaps also in response to heightened downregulation of the receptor.

### PAR1/4-deficient ECs have reduced PTP1B activity

We next sought to determine what drove the increased IR phosphorylation in PAR1/4-deficient ECs since this was likely responsible for heightened basal IR activity in these cells. We first looked at c-Src, a tyrosine kinase that can directly phosphorylate residues on the c-terminal loops of IR independently of ligand binding (23). However, we found that HUVECs treated with an inhibitor of c-Src (saracatinib) showed marginally elevated IR phosphorylation (∼1.28-fold, p<0.0766) (**Figure S13A&E**), rather than the decrease we expected, so we considered other modulators of IR activity.

Our next candidate was protein tyrosine phosphatase-1B (PTP1B/*Ptpn1*), which negatively regulates insulin signaling by dephosphorylating IR c-terminal residues (24). Recent studies have shown that PTP1B regulates IR activity in HUVECs (25). Furthermore, the inhibition of PTP1B results in enhanced insulin uptake in bovine aortic ECs (26, 27), suggesting a role for this receptor in endothelial insulin transcytosis. Lastly, PTP1B knockout mice show increased insulin sensitivity and glucose tolerance (28), although the endothelial contributions to these phenotypes are unclear.

Our analysis of *Ptp1b* transcripts in datasets generated from multiple mouse organs (29) revealed that the gene encoding this phosphatase is consistently enriched in ECs compared to total organ input samples (**Figure S15**), supporting its importance in ECs *in vivo* (30). Concordantly, we found that HUVECs treated with a PTP1B inhibitor (JTT-551) showed significantly increased IR phosphorylation (**Figure S13A&B)** and insulin transcytosis (**Figure S9D**) over control cells, like PAR1/4*-*deficient ECs.

To determine if loss of endothelial PARs affected PTP1B activity, we used a phosphotyrosine phosphatase activity assay (31) in conjunction with an allosteric inhibitor for PTP1B (32) to measure PTP1B activity in control and PAR1/4*-*deficient HUVECs. We validated this PTP1B inhibitor by demonstrating that it led to selective reduction of PTP1B activity while preserving non-PTP1B phosphatase activity in HUVECs (**Figure S16**). Using this approach, we were able to show that PTP1B activity constitutes (∼25.5%) of the total phosphotyrosine phosphatase activity in HUVECs (**Figure S16B**), which equates to ∼2.16 µU of PTP1B activity per microgram of protein (**Figure 6A**). Importantly, we found that PAR1/4-deficient

**Figure 6.**
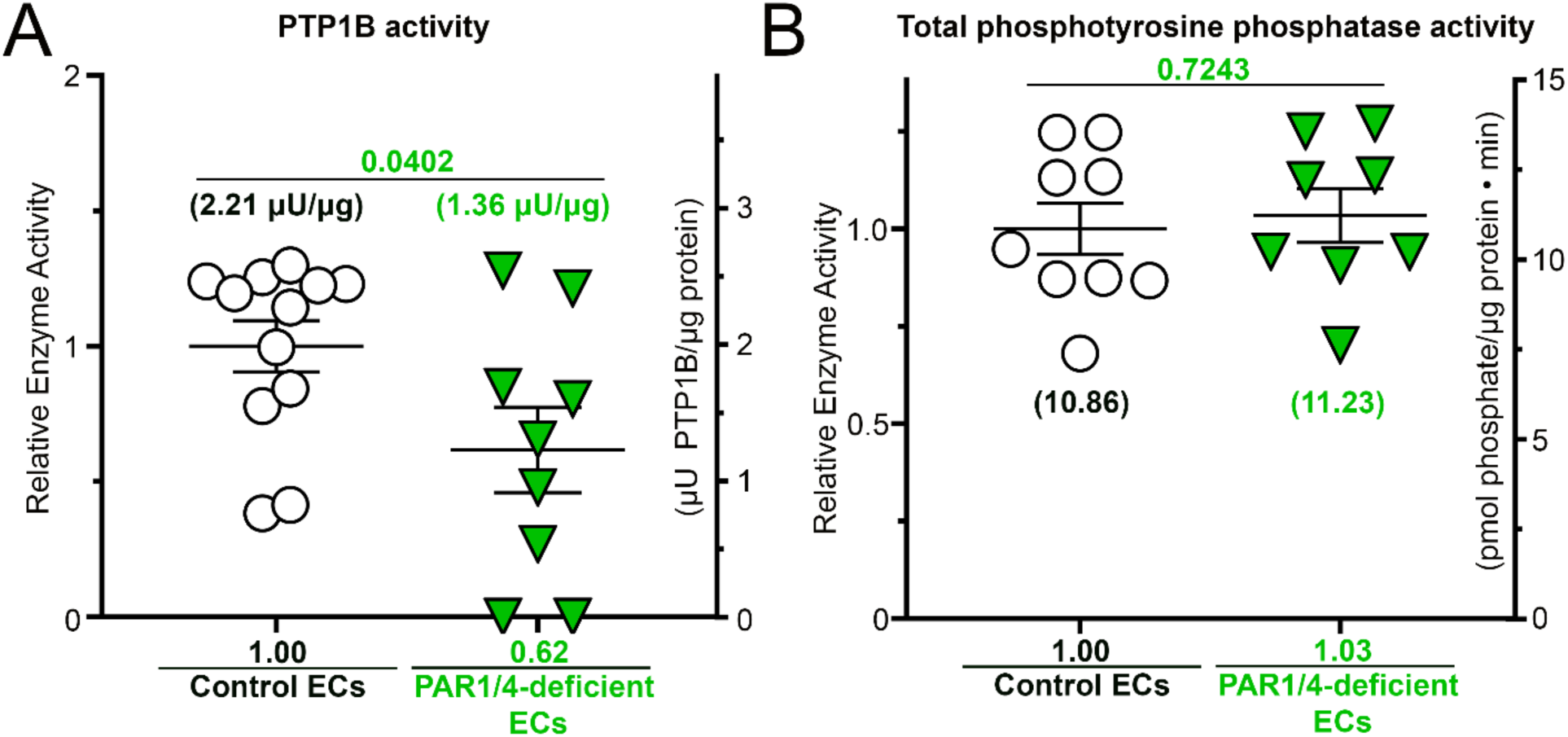
PAR1/4-deficient ECs have decreased PTP1B activity. (**A**) Relative PTP1B activity between control and PAR1/4-deficient HUVECs (n=9-12 technical replicates per group). (**B**) Relative phosphotyrosine activity between control and PAR1/4-deficient HUVECs (n=8-9 technical replicates per group). <u>Statistics</u>: (A-B) Analyzed by an unpaired t-test; p values are displayed in the graphs.

HUVECs had significantly reduced (∼37.0%; p<0.0402) PTP1B activity (∼1.36 µU per microgram of protein) (**Figure 6A**). This reduction in activity was not due to changes in PTP1B protein levels (**Figure S17**). Additionally, this reduction in PTP1B activity was selective since overall phosphotyrosine phosphatase activity was similar in control and PAR1/4-deficient ECs (**Figure 6B**). These data suggest that the loss of PAR1/4 in ECs reduces the activity of PTP1B, which may account for the rise in basal IR phosphorylation/signaling and insulin transcytosis we see in these cells.

This finding is particularly important since we also determined that STZ-induced diabetes significantly reduced *Insr* expression in quadriceps muscle ECs *in vivo* (∼62.5%, p<0.0411) (**Figure S18E**). This suggests that increasing endothelial IR activity may be beneficial in a diabetic context since the receptor’s expression is selectively reduced in ECs.

### Endothelial-specific deletion of one allele of *Insr* in *Par1/4^iECko^* mice restores STZ-induced hyperglycemia

To determine if the elevated IR activity we detected in PAR1/4-deficient ECs *in vitro* was related to the beneficial effects we saw on hyperglycemia in STZ-challenged *Par1/4^iECko^* mice, we used an *Insr*-floxed line to reduce one allele of *Insr* on ECs in the mutants (*Par1/4^iECko^;Insr^iEChet^*). We did not see significant differences between *Par1/4^iECko^* and *Par1/4^iECko^;Insr^iEChet^* mice when we challenged them at baseline with an acute ITT (**Figure S19**). However, when we challenged them with STZ, we found that STZ-treated *Par1/4^iECko^;Insr^iEChet^* mice showed higher blood glucose levels compared to STZ-treated *Par1/4^iECko^* mice one month after STZ treatment (**Figure 7A&B**). Altogether, STZ-treated *Par1/4^iECko^;Insr^iEChet^*mice showed significantly higher blood glucose AUC compared to STZ-treated *Par1/4^iECko^* mice (**Figure 7C**) (p<0.0438), with levels comparable to control mice treated with STZ. These results suggest that protection from STZ-induced hyperglycemia in *Par1/4^iECko^*mice is dependent on endothelial IR.

**Figure 7.**
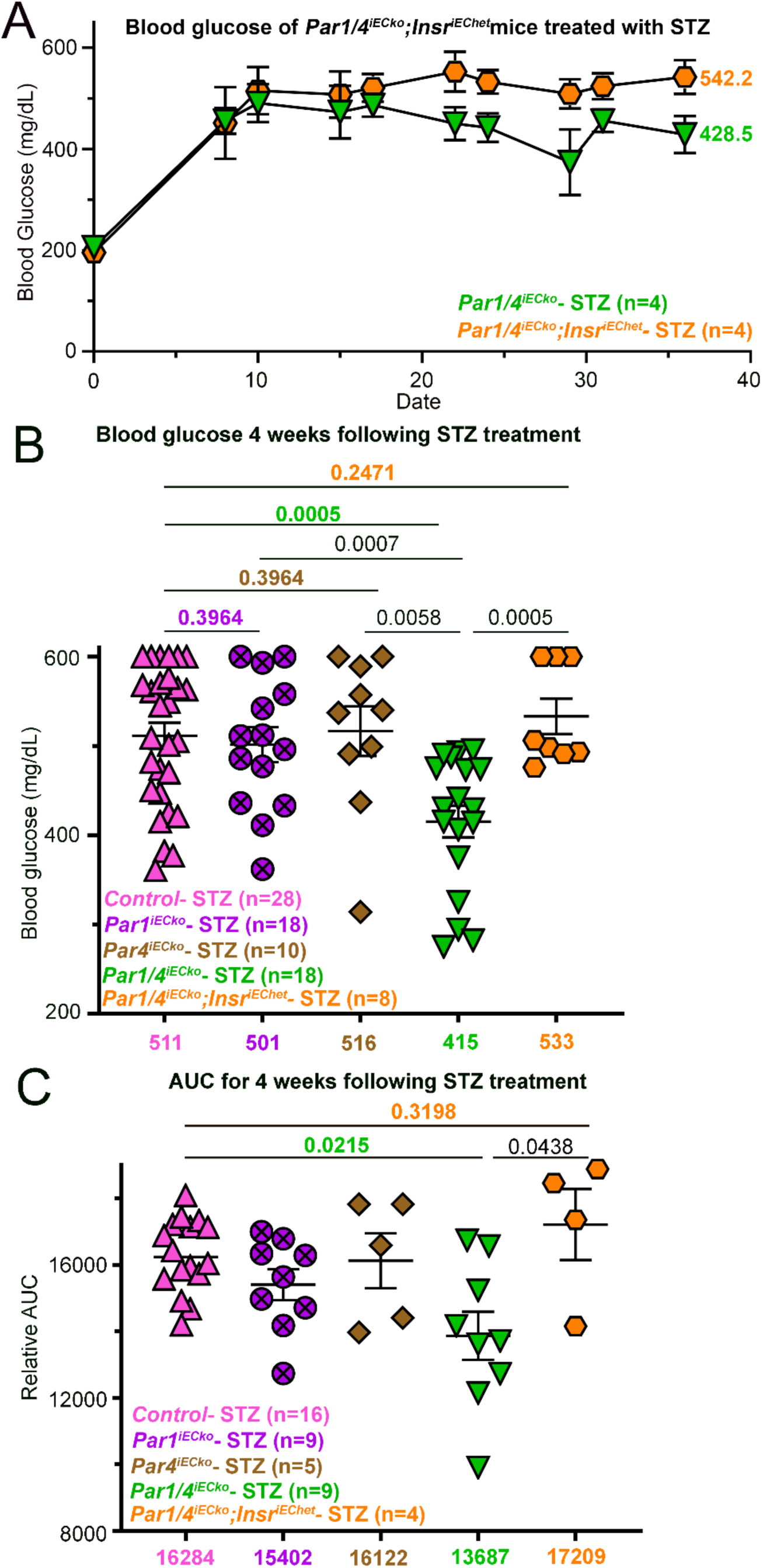
Endothelial-specific deletion of one allele of *Insr* in *Par1/4^iECko^* mice phenotypically restores STZ-induced hyperglycemia. **(A)** Blood glucose levels of STZ-treated male *Par1/4^iECko^* and *Par1/4^iECko^;Insr^iEChet^*mice. Diabetes was allowed to progress for 4 weeks. Final measurements taken at the end of the study are displayed on the graph. (n=4 mice per group). **(B)** Blood glucose levels were taken twice during the 4^th^ week of the study in A. (n=8-28 measurements from 4-14 mice per group). **(C)** Area under the curve (AUC) of blood glucose levels in A from the start of the study to 36 days (n=4-16 mice per group). <u>Statistics</u>: (B-C) Analyzed by Welch ANOVA followed by a 2-stage linear step-up procedure of Benjamini, Krieger, and Yekutieli for multiple comparison testing; q values are displayed in the graphs.

## DISCUSSION

The concept of endothelial insulin trafficking and transcytosis has been mired in controversy (4). As early as 1978, IRs were detected on cultured ECs (33), and within the next decade, insulin was shown to be rapidly internalized and transported through ECs with little to no degradation (34, 35). In mice, the presence of IR on small capillaries was shown to be critical for trafficking insulin to parenchymal tissues (36). From these studies, it was theorized that insulin is transcytosed across the vascular endothelium in an IR-dependent manner (4). However, later studies conducted in mice with EC-specific loss of IR (*Ir^ECko^*) showed limited changes in insulin sensitivity or glucose homeostasis, although they did show delays in insulin signaling in certain tissues (i.e., skeletal muscle, brown fat, and parts of the brain) (37, 38). This suggested a lesser role for endothelial IR in mediating insulin sensitivity. This latter finding may be related to insulin-like growth factor receptor 1 (IGF-1R), which is also present in the vascular endothelium and can mediate insulin transcytosis even when IR is absent in ECs (39, 40).

Even though there is mixed evidence that loss of IR reduces insulin sensitivity, there is ample evidence that increasing the activity of IR increases insulin uptake, transcytosis, and sensitivity in mice. Inhibition of PTP1B, a negative regulator of insulin signaling, results in increased IR activity and enhanced insulin uptake *in vitro* (25–27). Moreover, mice that display elevated levels of insulin receptor substrate 2 (IRS2)—a molecular adaptor of IR (15) whose elevation increases IR activity (15)—have increased insulin sensitivity. It is noteworthy that mice with EC-specific IRS2 deletion (*Irs2^ECko^*) display a greater reduction in insulin sensitivity than do *Ir^ECko^* mice (41, 42). However, given that IRS2 is recruited to both activated IR and IGF-1R in response to ligand stimulation of each receptor (43), this phenotype may mimic dual loss of endothelial IR and IGF-1.

Thus, genetic studies in mice suggest that modulating endothelial IR activity concordantly affects insulin sensitivity—particularly in cases of increasing IR activity. Our studies build on these findings by revealing a novel relationship between endothelial thrombin receptors (PAR1 and PAR4) and endothelial IR. We show that the loss of the endothelial thrombin receptors PAR1/PAR4 increases IR activity in ECs and whole-body insulin sensitivity in mice, thereby protecting them against STZ-induced hyperglycemia. Interestingly this effect requires dual loss of both PARs, although PAR1 deletion alone has a small but significant effect on STZ-induced hyperglycemia. These results are consistent with our previous study on PAR1/PAR4 in hepatic ECs, in which we showed these receptors act redundantly during the acute thrombin-generating challenge of acetaminophen overdose (2), despite their disparate expression levels (2, 3). In the present study, we also note that even without the STZ challenge, which modestly elevates circulating thrombin levels, *Par1/4^iECko^* mice have increased insulin sensitivity as measured by ITT and GTT. These data suggest that the whole-body insulin sensitivity phenotypes in STZ-challenged *Par1/4^iECko^*mice are driven by the loss of basal PAR signaling.

The elevated insulin sensitivity and IR activity we now report in PAR1/4-deficient ECs correlate with diminished PTP1B activity. Recent studies by Cho and colleagues (25) have shown a similar GPCR-PTP1B-IR signaling axis in ECs. They found that endothelial loss of the GPCR calcitonin receptor-like (CALCRL) results in reduced Gα_s_ signaling in ECs. This lower Gα_s_ signaling leads to reduced adenylate cyclase (AC) activity, reduced cyclic AMP (cAMP) levels, and reduced protein kinase A (PKA) activity. In ECs, PKA phosphorylates PTP1B on Ser_205_, which increases its activity and results in IR dephosphorylation. Thus, endothelial Gα_s_ signaling inversely correlates with IR phosphorylation and activity.

PARs are promiscuous GPCRs in that they couple to multiple heterotrimeric G proteins (Gα_12/13_, Gα_i/o_, Gα_q_) (3, 44). However, there is a notable lack of evidence for PARs coupling to Gα_s_. It is possible that Gα_s_ signaling could selectively be mediated by PAR1/4 heterodimers, which may account for why the dual loss of PAR1/4 is needed to drive IR activity. However, we previously demonstrated that PAR1/4 heterodimers only constitute a small proportion of PAR dimers in most tissues (3). It is also possible that PAR-mediated Gα_s_ signaling may require interaction with a secondary membrane protein, akin to the CALCRL-RAMP2 interaction needed for adrenomedullin signaling in ECs. Finally, it may be possible that our PAR-mediated effects on IR signaling are driven by a rise in Gα_i_ signaling, which reduces AC activity (45). We have previously shown that ECs display a homogenous composition of heterotrimeric G-proteins between different tissue beds, and the predominantly expressed endothelial G-protein is Gα_i_ (3). If the loss of endothelial PARs increases Gα_i_ signaling, this could mimic the loss of Gα_s_. Altogether, we acknowledge that more work will be needed to understand the mechanism by which PAR1/PAR4 deletion diminishes PTP1B activity and increases IR activity in ECs. For now, we know that the physical removal of PAR1 from ECs is required for our phenotypes because the inhibition of PAR1 with vorapaxar was not sufficient to increase IR activity in our studies.

We speculate that increased insulin transcytosis out of circulation and into parenchymal tissues drives insulin sensitivity in *Par1/4^iECko^* mice, but we acknowledge that other mechanisms may contribute as well. Previous studies have shown that in addition to insulin transcytosis, increased EC IR signaling also results in vasodilation and increased blood flow to target tissues by triggering the production of nitric oxide (NO) within ECs (25, 38, 41, 42, 46). This elevation in tissue perfusion increases glucose uptake. Therefore, it is possible that increased IR-mediated vasodilation also contributes to enhanced insulin sensitivity in *Par1/4^iECko^* mice. We also note that in diabetes there is a rise in thromboinflammation in tissues (47, 48), which leads to increased insulin resistance. We have shown that loss of endothelial PARs can reduce vascular permeability (2, 3); this may contribute to less thromboinflammation and protection against diabetes in *Par1/4^iECko^* mice. However, our genetic experiments involving the reduction of endothelial IR indicate that most of the beneficial phenotypes we see in STZ-challenged *Par1/4^iECko^* mice result from heightened IR activity in ECs. Nevertheless, PAR deficiency may mediate additional benefits beyond insulin sensitivity since *Par1/4^iECko^;Insr^EChet^* mice did not show different insulin sensitivities from *Par1/4^iECko^* mice in an acute ITT experiment.

Regardless of how the loss of endothelial PAR1/4 increases insulin sensitivity in mice, our studies demonstrate a relationship between these GPCRs and IR. However, there remains the open question of why such a signaling axis has evolved in ECs. Previous studies from our lab using kinome profiling predicted that PAR1 signaling in response to activated protein C (aPC) also increases IR activity in ECs (6). PAR1 mediates distinct signaling depending on the activating ligand: activation by aPC results in cytoprotective signaling characterized by anti-inflammatory/anti-apoptotic activity, whereas thrombin signaling results in pro-inflammatory/pro-apoptotic activity (6, 49). There are also reports that inhibition of basal PAR1 signaling can mediate anti-inflammatory effects (50), similar to aPC-mediated signaling. Why might aPC-activated PAR1 need to transactivate IR during anti-inflammatory/pro-survival signaling? The answer may be related to insulin roles in neural tissues. While insulin promotes glucose uptake in metabolic tissues, it primarily supports neuroprotection and pro-survival signaling in the brain and retina (51). We propose that aPC-activated PAR1 may hijack IR signaling to promote pro-survival responses in ECs. An unintended consequence of this relationship could be that the loss of endothelial PARs triggers a rise in endothelial IR activity, insulin transcytosis, and whole-body insulin sensitivity. Therefore, endothelial pro-survival signaling may inadvertently affect systemic metabolism. Altogether, our findings reveal a novel relationship between PARs and IR in ECs and identify a novel therapeutic axis for targeting diabetic pathology.

## LIMITATIONS OF STUDY

One limitation is that although *Par1/4^iECko^* mice were protected against STZ-induced hyperglycemia in this study, loss of PAR1 and PAR4 in ECs only marginally reduced hyperglycemia in mice with preexisting diabetes. We argue this represents an intrinsic limitation of the STZ model. The level of STZ-mediated islet death may be so high that the beneficial effects caused by the loss of endothelial PARs are not sufficient to overcome the diabetic burden. In the future, it would be interesting to test PAR deletion in a Type II diabetic model that is not associated with islet death. Another limitation related to the STZ model is that it does not fully capture the autoimmune component of Type I diabetes. However, since our study focused on how PARs affect global insulin sensitivity and trafficking, this model was sufficient to test roles for PARs in endothelial insulin signaling.

## MATERIALS AND METHODS

Detailed information on materials and methods used are provided in the online supplement.

## NON-STANDARD ABBREVIATIONS AND ACRONYMS

AUC: Area under the curve
BUN: blood urea nitrogen
ECs: Endothelial cells
GPCR: G Protein-Coupled Receptor
GSH: Glutathione
GST: Glutathione S-transferase
HUVECs: Human umbilical vein endothelial cells
IR: Insulin receptor
IGF-1R: Insulin-like growth factor 1 receptor
KBW: Kidney-to-body weight
MEC: Muscle EC
p85: Regulatory subunit of class IA PI 3-kinase
p110: p110 catalytic subunit of class IA PI 3-kinase
PAR1: Protease-activated receptor 1
PAR4: Protease-activated receptor 4
PI3K: Phosphoinositide 3-kinase
PTP1B: Protein tyrosine phosphatase 1B
TAT: Thrombin-antithrombin
STZ: Streptozotocin
TPM: Transcripts per million mapped reads
TRAP: Translating ribosome affinity purification

## ACKNOWLEDGEMENTS

The authors would like to thank Dr. Charmain Fernando Johnson (OMRF Cardiovascular Biology Research Program) for her helpful discussions during the preparation of this manuscript. Data processing and analysis were supported by the OMRF Center for Biomedical Data Sciences. Diagrams were created with BioRender.com.

## AUTHOR CONTRIBUTIONS

Conceptualization: RR

Formal analysis: RR

Funding acquisition: CTG, RR

Investigation: RR

Methodology: RR

Writing – original draft: RR

Writing – review & editing, CTG, RR

Visualization: RR

## DATA AND MATERIALS AVAILABILITY

The data reported in this article has been provided in the online supplement. All material will be shared upon reasonable request to the corresponding author.

