## Supplemental Methods, Figures, and Tables for "A novel function for endothelial protease-activated receptors in modulating insulin receptor activity with implications for diabetes"

### DETAILED MATERIALS AND METHODS

**Mice.** These studies included the following mice: *Par1*-flox (ViewSolid Biotech, Inc.) (1), *Par4*-flox (ViewSolid Biotech, Inc.) (1), *Cdh5(PAC)-Cre*$^{ERT2}$ (gift from Dr. Ralf Adams, Max Planck Institute for Molecular Biomedicine; also obtainable through Taconic: #13073) (2), *Rpl22*$^{tm.1.1Psam}$ (RiboTag; The Jackson Laboratory: #029977) (3), and *Insr*-flox (The Jackson Laboratories: #006955) (4). Inducible EC-specific deletion of *Par1, Par4,* and/or *Insr* was achieved by crossing mice expressing *Cdh5(PAC)-Cre*$^{ERT2}$ with *Par1*-flox and/or *Par4*-flox mice and/or *Insr*-flox mice, as described previously (1). For TRAP experiments, the RiboTag allele was homozygosed onto the *Par1/4*$^{iECko}$ line; RiboTag-*Par1*$^{iEChet}$;*Par4*$^{ECko}$ mice were used as controls for comparison. Genotyping of mutant alleles was conducted with the primers listed in Supplemental Table 1 (ST1). All mice were maintained on a C57Bl/6J background under environmental conditions that were previously described (1). All animal studies adhered to the ARRIVE guidelines proposed by the National Centre for the Replacement Refinement and Reduction of Animals in Research (NC3Rs). These guidelines include the design, conduct, interpretation, and reporting of animal studies. All experimental animal protocols were approved by the Institutional Animal Care and Use Committee at the Oklahoma Medical Research Foundation (Protocol: 21-28).

**Tamoxifen administration.** Gene deletion with the *Cdh5(PAC)-Cre*$^{ERT2}$ line was performed in 5–6-week-old mice with the administration of five doses of 10 mg/mL tamoxifen (200µL: 2 mg/day) (Sigma: #T5648) dissolved in peanut oil and administered via oral gavage every other day (total of 10 mg/mouse to achieve optimal gene deletion), as described previously (1). This approach was utilized since we have previously shown that loss of endothelial PAR1/4 is embryonic lethal (5); therefore, using a constitutively active endothelial Cre recombinase line would hinder the generation of experimental mice. Both Cre-positive and Cre-negative littermate controls were administered tamoxifen. Four weeks were allowed to pass between the last tamoxifen treatment and the first STZ treatment to prevent confounding results from the use of tamoxifen.

**Streptozotocin (STZ) administration.** STZ was diluted in 50 mM sodium citrate buffer, made by dissolving sodium citrate dihydrate (EMD Chemicals: SX0445-1) in autoclaved water. The buffer was filtered (0.22 um) and stored at -20°C. STZ was diluted in citrate buffer to 75 mg/mL, and 20 µL of this solution (1.5 mg) was administered to mice by intraperitoneal injection for 5 consecutive days totaling 7.5 mg/mouse. This corresponds to roughly 40-50 mg/kg/dose per mouse since the mice were weight matched at the start of the study. STZ injections occurred at 11 weeks of age, and mice were monitored for an additional 12-16 weeks. The study was terminated at 24 weeks for all mice.

**Sex as a biological variable.** Only male mice were used for the STZ studies because females metabolize STZ without pancreatic toxicity (6, 7). The numbers and sexes of all mice studied are listed in Supplemental Table 3 (ST3). To maintain consistency with the STZ studies, all other studies were conducted in male mice.

**Blood glucose monitoring via tail bleed.** Blood glucose was monitored to confirm induction of diabetes with STZ treatment and for ITT/GTT measurements. Animals were lifted out of their cage, placed in a work area, and gently restrained by holding the base of the tail. A sterile lancet was used to prick the tail. The tail was then gently squeezed from the base toward the tip to encourage blood flow. Approximately 0.5-1 µL of blood was collected for analysis with glucose test strips and a glucometer. Blood flow was then terminated by applying finger pressure to sterile gauze at the blood-sampling site (10-20 sec). The blood sampling site was monitored for infection. Blood glucose was monitored twice weekly during the STZ study.

**Mouse organ collection.** Mice were anesthetized with isoflurane using a Sigma Delta Vaporizer (Penlon #52606). Upon deep induction, mice were perfused, and organs were isolated as described previously with the following modifications (1). The thoracic cavity was opened to expose the heart and 300-500 μL of blood was isolated using a cardiac puncture extraction. A peristaltic pump (Bio-Rad Low-Pressure Chromatography Pump EP-1) was then used to run a perfusion solution (1X PBS supplemented with 1 mg/mL heparin sodium salt) for 10 min with a flow rate of 2 mL/min for a total of 20 mL of solution administered to each mouse. Following perfusion, kidneys were isolated and weighed upon isolation to quantify the kidney-to-body weight ratio. In the case of RiboTag mice, the perfusion was performed with a hand-operated syringe. 100 ug/mL cycloheximide (CHX) was supplemented to the perfusion solution and 50 mL was pumped into each mouse.

**Complete blood counts.** Before perfusion of the mouse, 300-500 μL of blood was collected by cardiac puncture. 150-250 μL of blood was placed in microtainer blood collection tubes containing $K_2$EDTA (Tiger Medical: TM28797). The tube was gently mixed and then analyzed using a Hemavet HV950FS (Drew Scientific) automated cell counter. The reference range provided in the figures are based on murine blood counts.

**Serum collection.** Before the mouse's perfusion, 300-500 μL of blood was collected by cardiac puncture. 150-250 μL of blood was placed in microtainer blood collection tubes containing a clot activator/SST™ Gel (Amber) (Tiger Medical: 365978). Blood was allowed to coagulate for 5 minutes and then centrifuged at 7,500 g for 5 min at 22°C. The serum was then isolated from the tube, flash-frozen in liquid nitrogen, and stored at -80°C for subsequent use.

**Renal toxicity assessment.** Mouse serum was subjected to albumin (ALB; IDEXX: 98-11065-01), blood urea nitrogen (BUN; IDEXX: 98-11070-01), and creatinine (CREA; IDEXX: 98-11074-01) measurements using a Catalyst One Chemistry Analyzer from IDEXX Laboratories, Inc. (Westbrook, ME).

**Plasma collection.** Before perfusion of the mouse 300-500 μL of blood was collected by cardiac puncture. 150-250 μL of blood was placed in microtainer blood collection tubes containing $K_2$EDTA (Tiger Medical: TM28797). The tube was gently mixed, and blood was allowed to sit at room temperature for 5 min to equilibrate. Blood was then centrifuged at 2,500 g for 10 min at 22°C. The plasma was then isolated from the tube, flash-frozen in liquid nitrogen, and stored at -80°C for subsequent use.

**Thrombin-antithrombin (TAT) ELISA.** Affinity-purified sheep anti-thrombin antibody (Affinity Biologicals SAFII-AP; 100 μL, 2 ug/mL) was incubated overnight on Immulon 4 HBX ELISA Plates (Thermo Scientific™ 3855). A BioRad Plate Washer (BioRad Model 1550) was used to wash the plate four times [Wash Buffer: 1X PBS (Thermo Scientific: 14190144), 0.05% Tween-20 (VWR: 0777-1L)] with 300 μL of wash buffer used per well for every wash. Plasma collected from mice was then diluted (1:100) in sample buffer [Sample Buffer: 1% BSA (Roche: 10775835001), 5 mM EDTA (Thermo Fisher: 15575020), 5 mM Benzamine Hydrochloride (Sigma: 63226-1mL-F), 0.1% Tween 20]. To each well of the plate, 20 μL of diluted sample was added and incubated for 2 hours under gentle rocking. The plate was then washed with the wash buffer 4 times. Affinity-purified biotinylated sheep anti-human antithrombin (ATIII) antibody (Affinity Biologicals SAAT-APBIO; 100 μL, 0.5 ug/mL) was diluted in a wash buffer and incubated for 1 hour. The plate was then washed again with the wash buffer another 4 times. Peroxidase streptavidin (Jackson Immuno: 016-030-084; 500 ng/mL) was diluted in 1X wash buffer and incubated for 30 minutes. The plate was then washed again with the wash buffer another 4 times. Finally, 100 μL of SuperSignal™ West Femto Maximum Sensitivity Substrate (Thermo Scientific: 37075) was then incubated on the plate, and luminescence was read immediately using a plate reader (BMG Labtech FlourStar Omega). TAT ELISA standards were made by incubating equivalent amounts of mouse thrombin (Molecular Innovations: MTHROM) and mouse antithrombin III (Haematologic Technologies: MCATIII-5120)—25 μM final concentration each in 1X PBS at 22°C for 5 minutes. The TAT solution (3.25 mg/mL) was then diluted to 5000 ng/mL and serially diluted to make a standard for the ELISA.

**Hyaluronan (HA) ELISA.** Isolated serum was assayed for hyaluronan levels by using a commercial kit (Echelon Biosciences; K-1200) following the manufacturer's instructions.

**Insulin ELISA.** Isolated serum was assayed for circulating insulin levels by using a commercial kit (ALPCO; K-80-INSHUU-E01.1) following the manufacturer's instructions. Serum was isolated from mice following 6 hours of fasting (*see serum collection*).

**Insulin tolerance test (ITT).** This test was employed to determine the sensitivity of insulin-responsive tissues in mice by measuring the glucose in circulation after a bolus intraperitoneal injection of insulin. One week before the experiment, mice were acclimated to handling by performing daily saline injections; this mitigates cortisol stress responses that temper the ITT results. Mice were fasted overnight before the ITT and provided water *ad libitum*. To match the results of our STZ study, only male mice were used for ITT testing. Mice were administered 1.0 IU insulin/kg body mass (Insulin: Novolin R U100). Blood glucose was then measured at 0 min, 15 min, 30 min, 60 min, 90 min, and 120 min (see "Blood glucose monitoring via tail bleed").

**Glucose tolerance test (GTT).** This test was employed to determine the sensitivity of glucose tolerance in mice (i.e., the clearance of glucose from the blood to tissues like muscle and fat) by measuring the glucose in circulation after a bolus intraperitoneal injection. One week before the GTT test, mice were acclimated to handling by performing daily saline injections; this mitigates cortisol stress responses that temper the GTT results. Mice were fasted overnight before the GTT and provided water *ad libitum*. To match the results of our STZ study, only male mice were used for GTT testing. Mice were then administered 1g glucose/kg body mass (Glucose: Sigma G8270). Blood glucose was measured at 0 min, 15 min, 30 min, 60 min, 90 min, and 120 min (see "Blood glucose monitoring via tail bleed").

**Reversed STZ experiment.** This study mimicked the STZ study shown above; however, in this case, we sought to make use of the inducible Cre to determine if the increased insulin sensitivity observed in *Par1/4*$^{iECko}$ mice would rescue hyperglycemia in mice treated with STZ. *Par1/4*$^{iECko}$ mice and littermate controls were treated with STZ at 6 weeks of age. STZ was diluted in citrate buffer to 75 mg/mL, and 20 µL of this solution (1.5 mg) was administered to mice by intraperitoneal injection for 5 consecutive days totaling 7.5 mg/mouse. Mice were then allowed to develop hyperglycemia over 4 weeks. On the 24$^{th}$ day, mice were administered tamoxifen—five doses of 10 mg/mL tamoxifen (200µL: 2 mg/day) dissolved in peanut oil via oral gavage every other day over two weeks (total of 10 mg/mouse). Blood glucose was then assayed in mice for 12 weeks to determine changes in hyperglycemia. The study was terminated at 20 weeks.

**Culture of human umbilical vein endothelial cells (HUVEC).** Primary HUVECs (ATCC, PCS-199-010) were grown in Endothelial Growth Media (EGM-2; Lonza CC-4176) media and serum starved in Endothelial Basal Media (EBM-2; Lonza CC-3156) media. Both these media were supplemented with 1% antibiotics [streptomycin/amphotericin (Sigma; #A5955)]. HUVECs were analyzed between passages 2 and 6 (P2-P6). Cells were split upon reaching 70% confluence in a plate. Cells were maintained in T-75 flasks (Cellstar: 658-170) and incubated in a cell culture incubator (Nuaire: NU-5810) at 5% $CO_2$. For thrombin studies, EGM-2 was replaced with EBM-2. Media was then supplemented with 50 mM glucose and/or 5U/mL thrombin. Cells were incubated with treatment for 12 hours before being isolated and sampled for changes.

**siRNA mediated gene knockdown of HUVECs.** For siRNA-mediated knockdown of PAR1, nonspecific (NS) control siRNA (Thermo; 4390844) or PAR1 siRNA (Thermo; 4390824-s4923) was transfected with Lipofectamine RNAiMAX reagent (Invitrogen; 56532). The siRNAs were prepared according to the manufacturer's instructions and were diluted alongside the Lipofectamine RNAiMAX reagent in Gibco™ Opti-MEM™ I Reduced Serum Medium (Fisher Scientific: 31-985-070). Lipofectamine-RNA complexes were incubated overnight, and the following morning the media was replaced with fresh EGM-2 media. Cells were collected for experimentation after another 24 hours. For transfection studies, cells were maintained in 100 mm cell culture dishes (Corning: 353003)

**Cell culture HEK-293.** HEK-293 cells were used to generate lentivirus. HEK-293 cells were grown in Dulbecco's Modified Eagle Medium (DMEM; Gibco; #11960-044) supplemented with 10% fetal bovine serum (FBS; Fisher Scientific; 3SH30910.03) and 1% antibiotics. Cells were split upon reaching 90% confluence in a plate. HEK-293 cells were used between P1 and P15. Cells were maintained in T-75 flasks (Cellstar: 658-170)

**Transcytosis assay.** The assay was modified from similar experiments conducted by Pillion and colleagues (8). HUVEC were seeded at (10,000-50,000 cells) on transwells with a membrane pore size of 0.4 μm (Costar 3460). Upon verification of confluency, HUVECs were washed and serum starved in EBM-2 media (Lonza CC-3156) for 2 hours. This step was performed to remove IGF-1 present in EGM-2 media (Lonza CC-4176) in which the cells were grown. The transwells were removed and added to a fresh plate containing no media in the bottom well; this was done to prevent paracellular leak in the subsequent step. Cells were then incubated with 500 nM biotinylated insulin (I2258, Sigma) diluted in EBM-2 media chilled to 4°C for 10 min. The media was chilled to allow the binding of biotinylated insulin to the cell surface but not internalization. The transwell was then washed with PBS to remove the unbound insulin. The transwells were then added to a fresh plate containing 1.5 mL of EBM-2 media. Pre-warmed EBM-2 media (37°C) was then added to the top transwells to initiate insulin internalization and release. After 30 min, the bottom chamber was sampled for biotinylated insulin using an ELISA.

**Biotinylated insulin ELISA.** Streptavidin-coated plates were made in-house by incubating 100 μL of 1 μg/mL streptavidin dissolved in 100 mM carbonate/bicarbonate buffer (pH 9.6) in Immulon 4 HBX ELISA Plates (Thermo Scientific: 3855) overnight at 4°C. Plates were then washed four times with a Wash Buffer [1X PBS (Thermo Scientific: 14190144), 0.05% Tween-20 (VWR: 0777-1L)] using a BioRad Plate Washer (BioRad Model 1550) using 300 μL per well. Biotinylated samples were then incubated on the streptavidin-coated plate for two hours alongside insulin standards. Plates were then washed four times with the wash buffer. The plate was then incubated with an anti-insulin antibody (BioRad: 7F8 (E6E5) for one hour. The plate was then washed four times with the wash buffer and incubated with an HRP-conjugated secondary antibody to mouse IgG (CST: 7076S) for one hour. The plate was washed again four more times with the wash buffer. TMB substrate (Thermo Scientific: N301) was then added to the well and the reaction was allowed to proceed for 5-15 minutes, followed by inactivation with 50 μL of 6N $H_2SO_4$ (Thermo Scientific: 035610.K2). Optical density at 450 nm was determined with a spectrophotometric plate reader.

**2-NBDG Uptake Assay.** The assay was modified from similar experiments conducted by Pillion and colleagues (9). HUVECs were plated in 12-well plates and cultured in EGM-2 media. Cells were treated with PAR1-siRNA (ThermoFisher Scientific: 4390824-s4923) 24 hours before experimentation. Treated cells were then washed with glucose-free DMEM (ThermoFisher Scientific: 11966025), and the EGM-2 media was replaced with glucose-free DMEM. ECs were then treated with the following: vehicle, 100 nM insulin (100 nM), LY294002 (CST: 9901S), HNMPA3-AM (Santa Cruz Biotechnology: sc-221730), and vorapaxar (MedChemExpress: HY-10119/C) for two hours. Following this the media was replaced with glucose-free DMEM supplemented with analog 2-[N-(7-nitrobenz-2-oxa1,3-diazol-4-yl) mino]-2-deoxy-d-glucose (2-NBDG) (100 µM) (ThermoFisher Scientific: N13195). The plates were then incubated at 37°C with 5% $CO_2$ for 30 min. Wells were then washed twice with pre-chilled 1X HBSS (ThermoFisher Scientific: 14025092) at 4°C. Cells in each well were then digested with trypsin (ThermoFisher Scientific: 25200056), pelleted, and resuspended in chilled 1X HBSS and maintained on ice for flow cytometry analysis. Cells were then strained through a 70-µm cell strainer (Fisher Scientific: 03-421-228) and resuspended in 2% FBS/HBSS after being pelleted. Propidium iodide staining solution (50 ng/mL) (BD Biosciences: 556463) was applied before analysis to identify dead cells. Cell events were collected using a BD FACSCelesta flow cytometer with 2 lasers (488 nm and 633 nm). Flow cytometry analysis was performed according to a standard procedure without permeabilization. All data were further analyzed and plotted with FlowJo software (BD Biosciences).

**DNA isolation.** For isolation of DNA of quantity up to 50 ug, QIAprep Spin Miniprep Kit (50) (Qiagen 27104) was used. Up to 20 mL of bacterial cultures were grown for a miniprep kit depending on high-copy/low-copy plasmids. Extraction was performed following the manufacturer's instructions, and DNA was eluted in a final volume of 30-50 µL. For isolating DNA of quantity up to 1 mg, QIAGEN Plasmid Maxi kits were used (Qiagen: 12165) were used. Up to 500 mL of bacterial cultures were grown for a miniprep kit depending on high-copy/low-copy plasmids. Extraction was performed following the manufacturer's instructions, and DNA was eluted in a final volume of 200 µL.

**Cloning.** pSicoR constructs originating from Addgene (Plasmid #11579) were cloned to include the addition of a PAR1-shRNA sequence and/or a Blasticidin resistance cassette. This cloning was performed using a Gibson assembly kit from Takara (Takara Bio: 638945). InFusion cloning was performed following the manufacturer's instructions.

**Viral transduction of HUVECs.** *Day 1:* HEK-293 cells (<P15) (ATCC: CRL-1573) were plated on a 100 mm tissue culture plate (VWR: 734-2341) with DMEM supplemented with 10% FBS. *Day 2:* a vial of HUVECs containing (50,000 cells) was thawed and plated in a T25 flask (ThermoFisher Scientific: 156367). On the same day, the plate containing HEK-293 cells was transfected with lentivirus [Lentivirus: psPAX2 (6 µg), PMD2.G (1.5 µg), Transfer Plasmid (7.5 µg)]. This cocktail was dissolved in Gibco™ Opti-MEM™ I Reduced Serum Medium and packaged with Lipofectamine 2000 (ThermoFisher Scientific: 11668019). DNA cocktail was diluted in 300 µL Opti-MEM and 50 µL Lipofectamine 2000 was diluted in 700 µL Opti-MEM. The two dilutions containing DNA were mixed and allowed to incubate at room temperature for 5 minutes. DMEM was removed from the plate of HEK cells and was replaced with EGM-2 media. The transfection mixture was then added dropwise to the plate of HEK cells. *Day 3:* EGM-2 media was replaced in the HEK-293 cell plates, with fresh EGM-2 media supplemented with 10 µg/mL polybrene (Sigma: TR-1003-G). On the same day, HUVECs were split into multiple T-25 flasks with a target confluency of 20-30%. *Day 5:* EGM-2 media was replaced in HEK-293 cell plates, with fresh EGM-2 media supplemented with 10 µg/mL polybrene (Sigma: TR-1003-G). 9 mL collected media was centrifuged at 2000 X *g* for 10 minutes to pellet dead cells. The collected media was then filtered through a 0.2 uM syringe filter (VWR: 28145-477) and added to the flask of HUVECs that had

been split on day 3. 1 mL of fresh EGM-2 media supplemented with 10 µg/mL polybrene was added to the flask. Plates were imaged using the ZOE fluorescence imager to verify transfection efficiency. *Day 7:* EGM-2 media was replaced in HEK-293 cell plates. 9 mL collected media was centrifuged at 2000 X *g* for 10 minutes to pellet dead cells. The collected media was then filtered through a 0.2 uM syringe filter (VWR: 28145-477) and added to the flask of HUVECs. 1 mL of fresh EGM-2 media supplemented with 10 µg/mL polybrene was added to the flask. Plates were imaged using the ZOE fluorescence imager to verify viral transduction efficiency. *For transfer plasmid containing blasticidin resistance cassettes for cell selections; Day 10:* EGM-2 media was removed and replaced with EGM-2 media supplemented with 5 ug/mL blasticidin S HCl (Gibco: A11139-03). This media was replaced on Day 13 with EGM-2 media supplemented with 5 ug/mL. By *Day 15* death of non-transduced cells had occurred. Media in plates was replaced with EGM-2 media, and the cells were allowed to proliferate and used for subsequent experiments. Cells were imaged using a ZOE fluorescence imager to verify viral transduction efficiency and blasticidin selection.

**Microscopy and image acquisition.** HUVEC and HEK-293 cells were imaged using a ZOE Fluorescent Cell Imager (BioRad: 1450031) for verification of viral infection. Confocal imaging was performed using an Olympus 1200 confocal microscope.

**Immunoblot**. Frozen mouse tissue samples or cells were first lysed in 3T3 buffer [1% IGEPAL (Sigma 56741), 20 mM HEPES (Sigma H3375) pH 7.4, 2 mM EDTA (Thermo Fisher: 15575020), 100 mM sodium fluoride (Sigma S6521), 10 mM sodium pyrophosphate (Sigma 221363), 1 mM sodium orthovanadate (MP Biomedical 159664), 1 mM PMSF (Sigma P7626), 1 mM sodium molybdate (Sigma M1003) and a protease inhibitor tablet (Roche: 11873580001)]. Tissue samples were pulverized using a mini bead beater (DENTSPLY; #C321001). Lysate was then sonicated on ice with 3 pulses with an amplitude of 20 units. Each pulse lasted 20 seconds, with a 40-second pause between pulses. Lysate concentrations of protein were then measured with a BCA Protein Assay Kit (Thermo Scientific; #23227) following the manufacturer's instructions. Samples were diluted to a concentration of 1 µg/µL through the addition of 3T3 lysis buffer and 2X Laemmli buffer (4% SDS, 20% glycerol, 0.004% bromphenol blue, 0.125M Tris-HCl (Roche, 10812846001), pH 6.8, 10% 2-mercaptoethanol). The samples were then boiled at 100°C for 5 minutes. Protein samples were then run on a 10% SDS-PAGE protein gel consisting of a loading gel [1.5 M Tris-HCl pH 8.8, 0.4% SDS, 10% Acrylamide (BioRad 1610158), 0.00027% Ammonium Persulfate (APS; Sigma 215589), 0.0013% TEMED (Santa Cruz sc-29111)] and a stacking gel (0.5 M Tris-HCl pH 6.8, 0.4% SDS, 7.5% Acrylamide, 0.00070% APS, 0.0035% TEMED), which were cast in 1.5 mm cassettes (Thermo Scientific: NC2015) and then transferred to a nitrocellulose membrane using iBlot Transfer Stacks (Thermo Scientific; #IB23001). The membranes were then blocked in 5% milk (BioRad: 1706404) or 5% BSA (for probing of phospho-proteins) dissolved in modified TBST (20 mM Tris-HCl pH 7.4, 400 mM NaCl, 0.001% Tween-20) and incubated in primary antibodies overnight at 4°C. The following day the blots were washed three times with TBST; each wash was 15 minutes. After these washes, secondary antibodies were applied for 1 hour at room temperature followed by another three washes with TBST. Chemiluminescence was performed to visualize bound antibodies; West Dura (Thermo Scientific 34076) and West Femto (Thermo Scientific 34095) were used. Membranes were imaged using an iBright CL750 Imaging System (Thermo Scientific; #A44116). NIH ImageJ software was used to quantify band density.

**p85-GST probe generation.** p85-GST probes were purified as described previously (10) with the following modifications. Fusion vectors containing GST and GST fused to the p85 subunit of PI3K (GST-p85) were transformed into BL21 (DE3) maximum efficiency *E. coli* competent cells (Thermo Fisher: EC0114) for protein purification. BL21 cultures containing the fusion vector were plated on Terrific Broth (ThermoFisher Scientific: 22711022) (TB)-Carbenicillin (Sigma: C3416) (100 µg/mL) plates and grown overnight. Individual clones were picked and inoculated in a liquid culture of 50 mL TB/Carb (100 μg/mL), and these cultures were grown overnight. The following morning these cultures were diluted in TB/Carb (1:10) medium grow for 2 h at 37°C. GST protein synthesis was initiated with the addition of isopropyl-B-D-thiogalactoside (IPTG) (Roche: 11422446001) to 0.1 mM final concentration for 1 h at 37°C. Bacterial cultures were collected and spun at 3000 × *g* for 10 min at 4°C, and the medium was decanted. The pellets were resuspended and washed in 10 mL ice-cold PBS. The suspension was then spun at 12,000 × *g* for 10 min at 4°C. The PBS was decanted, and pellets were resuspended in 1 mL lysis buffer [50 mM HEPES (pH 7.5), 50 mM NaCl, 10% glycerol, 1% Triton X-100 (ThermoScientific: A16046.AE), 1 mM EDTA, 1mM EGTA (EMD Millipore: 324626), and 1 mM phenylmethylsulphonyl fluoride (PMSF), supplemented with a protease inhibitor tablet (Roche: 11873580001)]. Lysate was then sonicated on ice with 3 pulses with an amplitude of 50 units. Each pulse lasted 10 seconds, with a 30-second pause between pulses. The lysate was then spun at 16,100 × *g* for 10 min at 4°C, and the supernatant was collected. 300-500 μL of Glutathione Sepharose (GSH) (50% v/w) beads were added to the lysate and rotated for 30 min at 4°C. Beads were then spun down at 16,100 × *g* for 1 min at 4°C and resuspended with 1 mL 1X PBS. Beads were then spun and resuspended two more times with 1 mL of 1X PBS. Beads were spun down a final time and resuspended in 500 μL lysis buffer. To verify the binding of the fusion protein to the GSH beads, 5-20 μL of beads were run on an SDS-PAGE gel. Beads were added to an equal volume of 2X Laemmli buffer, incubated for 5 min at 100°C, and resolved on a 10% SDS-PAGE gel. The gels were stained with Imperial Protein Stain (ThermoScientific: 24615). The gels were washed with boiling water, three times for 15 minutes each. The gel was then allowed to stain for 1 hour with imperial stain with gentle shaking. The gel was then destained in water overnight. SDS-PAGE gels of GSH bead-bound fusions were transferred to a nitrocellulose membrane using iBlot transfer stacks and probed with a GST antibody (*see immunoblot*).

**p85-GST pulldown.** Pulldowns were performed as described previously (10) with the following modifications. To assess insulin receptor activity/phosphorylation, a pulldown was performed with the p85-GST probes. 25-50 µL of Glutathione Sepharose 4B beads bound to p85-GST as well as Glutathione Sepharose 4B beads bound to GST alone (5-10 µL) (control) were added to 75–250 µg of tissue or cells which were lysed in 3T3 buffer (*see immunoblot*). The reactions rotated at 4°C for 2 hours. Beads were then spun down at 16,100 × g for 1 min at 4°C and resuspended in 500 µL wash buffer [50 mM HEPES (pH 7.4), 118 mM NaCl, 100 mM NaF, 2 mM Sodium orthovanadate, 0.1% (w/v) SDS, and 1% (v/v) Triton X-100]. Beads were then spun down at 16,100 × g for 1 min at 4°C. GST fusions were washed two more times with wash buffer. 25 µL 2X Laemmli buffer was then added to beads, and samples were boiled for 5 min at 100°C. The proteins were then resolved on a 10% SDS-PAGE gel, then transferred to a nitrocellulose membrane using an iBlot transfer system and subsequently probed with an anti-IRβ antibody (CST: 23413S)

**<u>Endothelial Translating Ribosome Affinity Purification (TRAP).</u>** We used a similar method for TRAP that we previously described (1). Frozen quadricep muscles (25-100 mg) were pulverized using a BioPulverizer (Biospec Products: 59012N), with 750 µL of polysome buffer [50 mM Nuclease free Tris pH 7.5 (BioWorld: 21420063-3), 1% Igepal CA-630 (Sigma: 56741), 1 mM dithiothreitol (Sigma: 10197777001), 100 µg/mL cycloheximide (Sigma: 01810), 100 mM KCl (Sigma: P3911), 12 mM $MgCl_2$ (Sigma: M8266), 1 mg/mL heparin sodium salt (Sigma: H3393), 200 U/mL RNaseOUT (Thermo Fisher: 10777019), and an EDTA-free protease inhibitor cocktail (Sigma: 11836170001), polysome buffer was prepared in DEPC-treated nuclease-free water (Thermo Fisher: AM9906)]. Tissue homogenates were then centrifuged twice at 16,000 g at 4°C for 15 min. The supernatant was isolated and diluted to 600 µl with polysome buffer. 100 µl of homogenate was aliquoted to form the input fraction, against which the EC-enriched samples could be compared. From the remaining homogenate, 400 µL was isolated and incubated with 5 µL (1:80) purified monoclonal rabbit anti-HA antibody (CST: C29F4) for 1 hour at 4°C under gentle rocking. 100 µL of magnetic protein G beads (New England BioLabs: S1430S) were then added to the antibody-incubated homogenates for an additional 30 min at 4°C under gentle rocking. High salt buffer (polysome buffer with the KCl concentration adjusted to 300 mM) was used to wash these magnetic beads three times. 500 μL of Trizol/chloroform was added to the beads to extract RNA from the bound polyribosome and the input fractions. Samples were centrifuged at 16,000 x *g*, and the upper layer was added to an equal volume of 70% ethanol. RNA was isolated from this solution using a Qiagen RNeasy Mini Kit, including an on-column DNase digestion step. RNA was eluted with 30 μL of DEPC-treated nuclease-free water and concentrations were quantified spectrophotometrically with a NanoDrop.

**PTP1B activity assay.** PTP1B activity was measured as previously described (11) with the following modifications. HUVECs were cultured as described above in 100 mm dishes. Cells were treated with siRNA for PAR1 and collected 48 hours following treatment. The cells were then lysed in a custom lysis buffer [100 mM HEPES, 2 mM EDTA, 150 mM NaCl, 2 mM PMSF, 0.1 mM Leupeptin (Sigma: L2884), 1% IGEPAL, and 2 μg/mL N-acetyl-L-leucyl-L-leucyl-L-norleucinal (Sigma: 11086090001)]. The cells were lysed in a volume of 100 μL yielding an approximate protein concentration of 2-4 μg/μL, which was quantified using a BCA assay. EC lysates were then diluted (1:2) in assay buffer [100 mM HEPES, 2 mM EDTA, 1 mM DTT (Sigma: 10197777001), 150 mM NaCl, and 0.5 mg/mL BSA]. Reactions were prepared in 35 μL total volume. Each sample was prepared in duplicate, one which was prepared in assay buffer and the other which was prepared in assay buffer that was supplemented with an allosteric PTP1B inhibitor (EMD Millipore: 539741-5MG) (12). 100 μM of this inhibitor was used in the relevant reactions. These reactions were then kept at 30°C for 15 minutes. This was performed since tyrosine phosphatases have high binding affinity for their cognate peptides; to ensure adequate inhibition of the enzyme this preincubation step was performed to give the inhibitor time to bind and inactive PTP1B. To these reactions, a phosphatase substrate (Echelon Biosciences: 871-70) (Sequence: TEGQ(pY)QPQP-OH) at a final concentration of 125 μM was added per reaction. This reaction was then allowed to progress at 30°C for 30 minutes. During this time, protein tyrosine phosphatases liberated the inorganic phosphate from the tyrosine residue of the substrate. After this incubation, 100 μL of malachite green solution

(Echelon Biosciences: K-1501) was added to each well. The liberated phosphate, which correlates to the phosphotyrosine phosphatase activity of each sample, formed a blue-colored complex with molybdate/malachite green. This color was allowed to develop at room temperature for 3 minutes and was quantified by reading the absorbance at 620 nm. Phosphate levels were normalized using a standard of known concentrations of inorganic phosphate (4000 picomoles – 0 picomoles) (Upstate Cell Signaling Solutions: 20-103). We measured the amount of liberated phosphate between our control and our PAR1/4-deficient ECs and were able to calculate the activities of phosphotyrosine phosphatases between the two groups. To determine changes in PTP1B activity, we measured the difference of liberated phosphate between our control and our PAR1/4-deficient ECs and their respective samples that were pretreated with a PTP1B-specific inhibitor. Using this differential, we could indirectly calculate the PTP1B activity between each group. To quantify the PTP1B activity between each group in terms of units, we measured the activity of pure PTP1B enzyme (EMD Millipore: KP8401-5UG). Using this we were able to determine that PTP1B had a specific activity of 108 $\mu mol \cdot min^{-1}\ mg^{-1}$. Using this value, we calculated the relative PTP1B activity in our ECs lysates.

**Immunocytochemistry.** Coverslips (FisherScientific: 50-949-314) were autoclave sterilized and coated in a collagen (Sigma: C4243) solution for 30 minutes at 37°C in the cell culture incubator (Nuaire: NU-5810). Coverslips were then washed in 1X HBSS and plated in the bottom of a 24-well plate (Cellstar: 662160). Cells were then seeded at low confluency (25,000 cells) in each well. 48 hours later cells were treated with a PAR1-siRNA (see *siRNA-mediated gene knockdown of HUVECs*). The following day media was replaced in the wells. 24 hours later the coverslips were removed, washed once with 1X HBSS, and fixed in 4% PFA [16% PFA stock (Electron Microscopy Science: 15710) diluted in 1X HBSS] for 15 minutes at room temperature. Cells were then blocked with 10% FBS for 1 hour at room temperature. FBS was diluted in 1X PBS supplemented with 0.1% Triton X-100 (PBS-T). Cells were then washed once with PBS-T and incubated with anti-IRβ antibody (CST: 23413S) diluted (1:25) in PBS-T and incubated overnight at 4°C. Coverslips were then washed once for 5 minutes with PBS-T and incubated at room temperature for one hour with a donkey anti-rabbit FITC conjugated antibody (ThermoFisher Scientific: A-21206) diluted to 1 μg/mL. Coverslips were then washed three times with PBS-T and incubated with DAPI (Sigma: D9542) at a concentration of (1:1000) for 10 minutes. Coverslips were then washed three more times with PBS-T and fixed in 50% glycerol. Cells were then imaged using an Olympus FV1200 confocal imager.

**Real-Time Quantitative Reverse Transcription PCR (qPCR)**. iScript cDNA Synthesis Kit (BioRad: 1708890) was used to convert RNA to cDNA (100-1000 ng) in a 20 μL reaction volume. Samples were incubated in the thermal cycler as per the manufacturer's instructions. Prepared cDNA was then diluted in DEPC-treated nuclease-free water to a final concentration of 3ng/5μL. SsoAdvance Universal SYBR Green Supermix (BioRad: 1725275) was then used to perform qPCR as per the manufacturer's instructions. To quantify the expression of IR type A (*INSR-A/Insr-A*), a splice primer was used to quantify the expression level of the type B spliceform. The result was normalized to total *INSR*/*Insr*, and then the reciprocal of that value was taken to give an indirect measure of the expression of *INSR-A/Insr-A*. The expression of type A was then normalized to the control sample.

**Structural modeling.** To show the structure of insulin receptor type A and type B (IR-A and IR-B, respectively), we generated *in silico* models of the receptors using RaptorX (13) (http://raptorx6.uchicago.edu/), as described previously (1). IR-A and IR-B are distinguished by a 12-residue peptide found in two of four insulin binding sites for the receptor. IR-A lacks this 12-residue peptide, and IR-B contains this peptide. To generate isoform-specific models we modeled two 220-residue fragments: one that contained this peptide and one that lacked this 12-residue peptide. These fragments were overlaid onto the existing Cryo-EM model of IR (6PXV) by Uchikawa, et al. (14), which had these fragments removed. Structural data were analyzed by UCSF Chimera (1.17.3) (15-17) (https://www.cgl.ucsf.edu/chimera/) and PyMOL (4.6.0). Autodock Vina (1.2.0) (18, 19) (https://vina.scripps.edu/) was used for *in silico* docking studies between insulin and the insulin binding sites of IR-A and IR-B. Docking studies were performed as previously described (20).

**Statistical analysis.** All statistical analyses were conducted as described previously (1). Analysis was performed using GraphPad Prism version 10.2.3. All scatter plots graphs display results as mean ± SEM. Each dataset includes a minimum sample size of (n=3) biological replicates, which was sufficient for all inferential testing.  Data analysis followed previously published criteria (1).

**Testing between two groups:** An unpaired t-test was performed when analyzing data that included one independent variable, normal distribution, and equal variances. Welch's t-test was performed when analyzing data that included one independent variable, normal distribution, and unequal variances. Mann Whitney U-test was performed when analyzing data that included one independent variable and non-normally distributed data.  The Wilcoxon Matched-Pairs Signed-Rank Test was performed when analyzing paired data that included one independent variable and non-normally distributed data.

**Testing between multiple groups:** One-way ANOVA was used for data that included one independent variable, and three or more independent groups with equal variances as measured by an F-test. Two-way ANOVA was used for data that included two independent variables and four groups with equal variances as measured by Spearman's rank correlation test to detect the presence of heteroscedasticity. Three-way ANOVA was performed for data that included three independent variables and eight groups with equal variances as measured by Spearman's rank correlation test to detect the presence of heteroscedasticity. Welch's ANOVA was performed on data that included one independent variable, and three or more independent groups that did not have equal variances, as determined by either a Spearman's rank correlation test to detect the presence of heteroscedasticity or an F test. The Friedman Test was performed on paired data that was normally distributed which included one independent variable, and three or more independent groups that did not have equal variances, as determined by an F test.

**Post hoc testing:** Testing was conducted using a two-stage step-up method by Benjamini, Krieger, and Yekutieli. This post-hoc test was chosen since it is less conservative than Tukey but still has significant power due to controlling the false discovery rate. When this method was performed q values are displayed in the figure. Note that tests for normality (Shapiro-Wilk) and equal variance (Spearman or F Test) were performed on all data. D'Agostino & Pearson normality tests were performed for samples with sufficiently large sample sizes. Two- and three-way ANOVAs were still performed even when a single data point caused the normality test and/or equal variance test to fail since these ANOVAs can still generate valid outcomes if the dependent variables are approximately normally distributed for the compared groups according to the central limit theorem. However, samples with different variances employed Welch's ANOVA to determine significant differences. A summary of all the statistical tests plus the n numbers used in all the figures can be found in Supplemental Table 3 (ST3).

**SUPPLEMENTAL FIGURES**

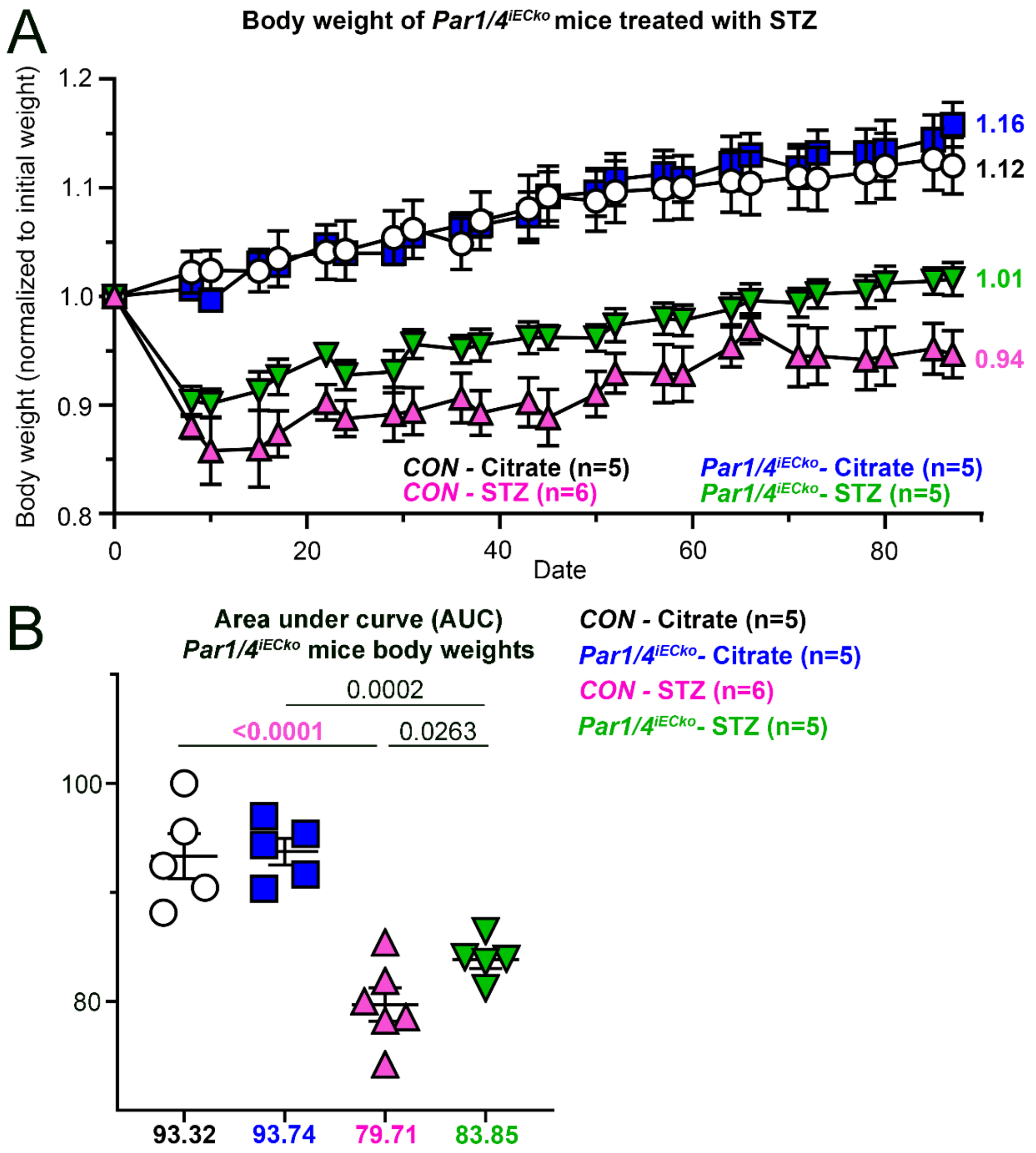

**<u>Figure S1.</u>** ***Par1/4*$^{iECko}$ mice show attenuation in weight loss following STZ induction. (A)** Body weights were measured in *Par1/4*$^{iECko}$ and littermate controls (CON) following treatment with STZ. Weights were normalized to the initial body weight prior to STZ treatment. **(B)** Area under the curve (AUC) for the upper panel. Analyzed by two-way ANOVA followed by a 2-stage linear step-up procedure of Benjamini, Krieger, and Yekutieli for multiple comparison testing; q values are displayed in the graphs. All scatter dot plots show mean ± SEM.

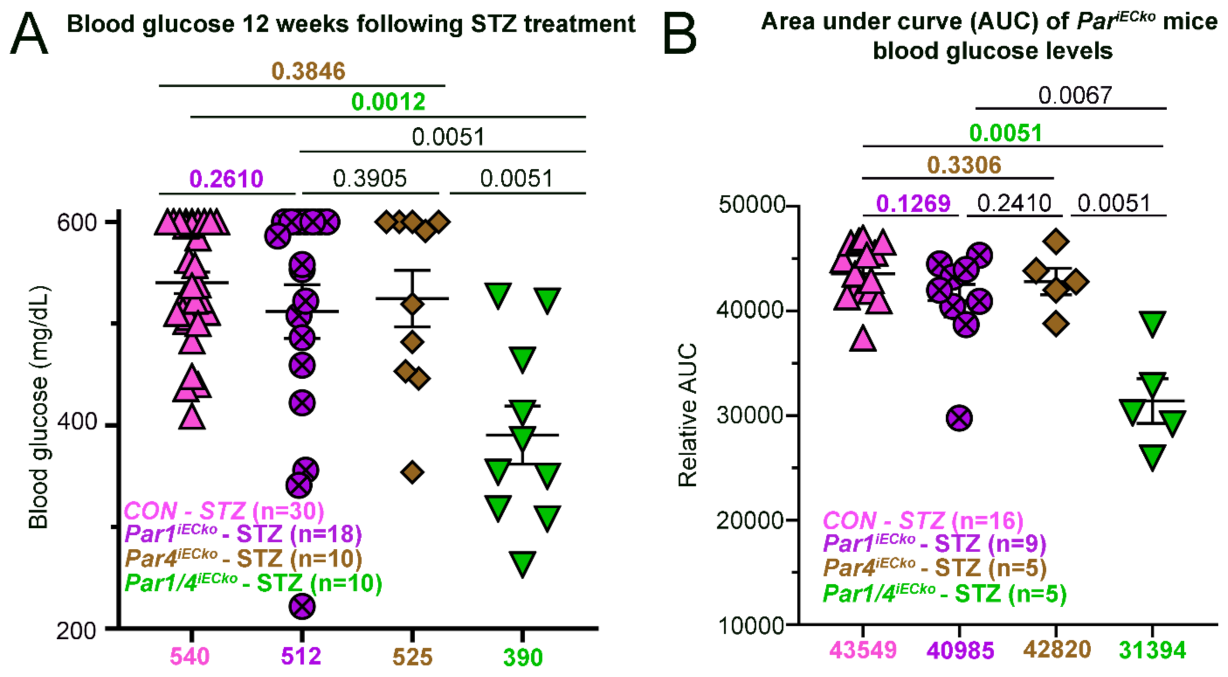

**<u>Figure S2.</u> STZ-treated $Par1/4^{iECko}$ mice display reduced blood glucose compared to control, $Par1^{iECko}$, and $Par4^{iECko}$ mice. (A)** Blood glucose levels were taken twice from $Par^{iECko}$ mice during the 12th week of the study and are graphically summarized above. (n=10-30 measurements from 5-15 mice per group) **(B)** Area under the curve (AUC) values of hyperglycemia for $Par1/4^{iECko}$ mice and littermate controls (CON) treated with citrate and STZ. <u>Statistics</u>: (A-B) Analyzed by Welch ANOVA followed by a 2-stage linear step-up procedure of Benjamini, Krieger, and Yekutieli for multiple comparison testing; q values are displayed in the graphs. All scatter dot plots show mean ± SEM.

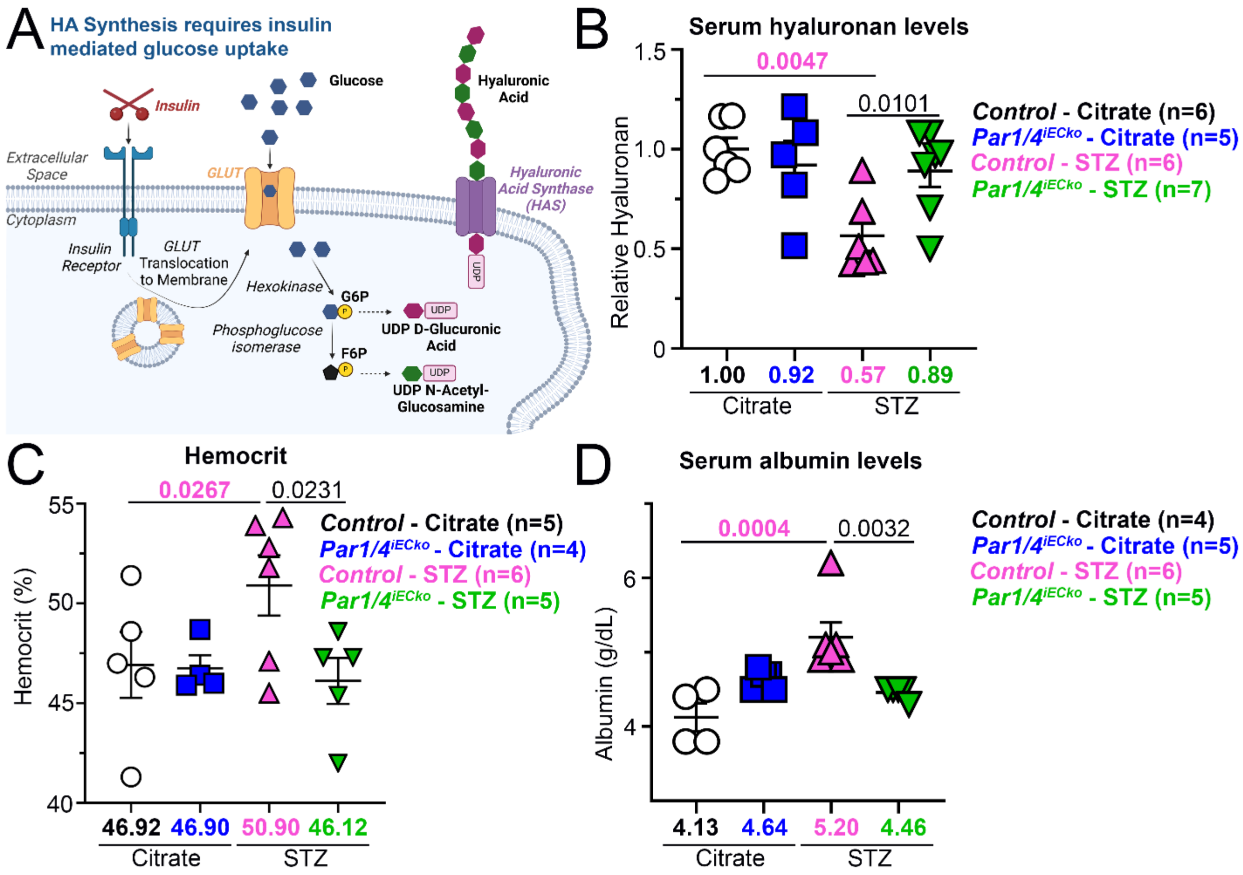

**Figure S3. STZ-treated *Par1/4*$^{iECko}$ mice display a reduction of hyperglycemic pathologies (A)** Schematic of how glucose drives hyaluronan synthesis. (**B**) Serum hyaluronan, (**C**) hematocrit, and (**D**) albumin were measured in mice after 12 weeks of STZ treatment. Analyzed by two-way ANOVA followed by a 2-stage linear step-up procedure of Benjamini, Krieger, and Yekutieli for multiple comparison testing; q values are displayed in the graphs. All scatter dot plots show mean ± SEM.

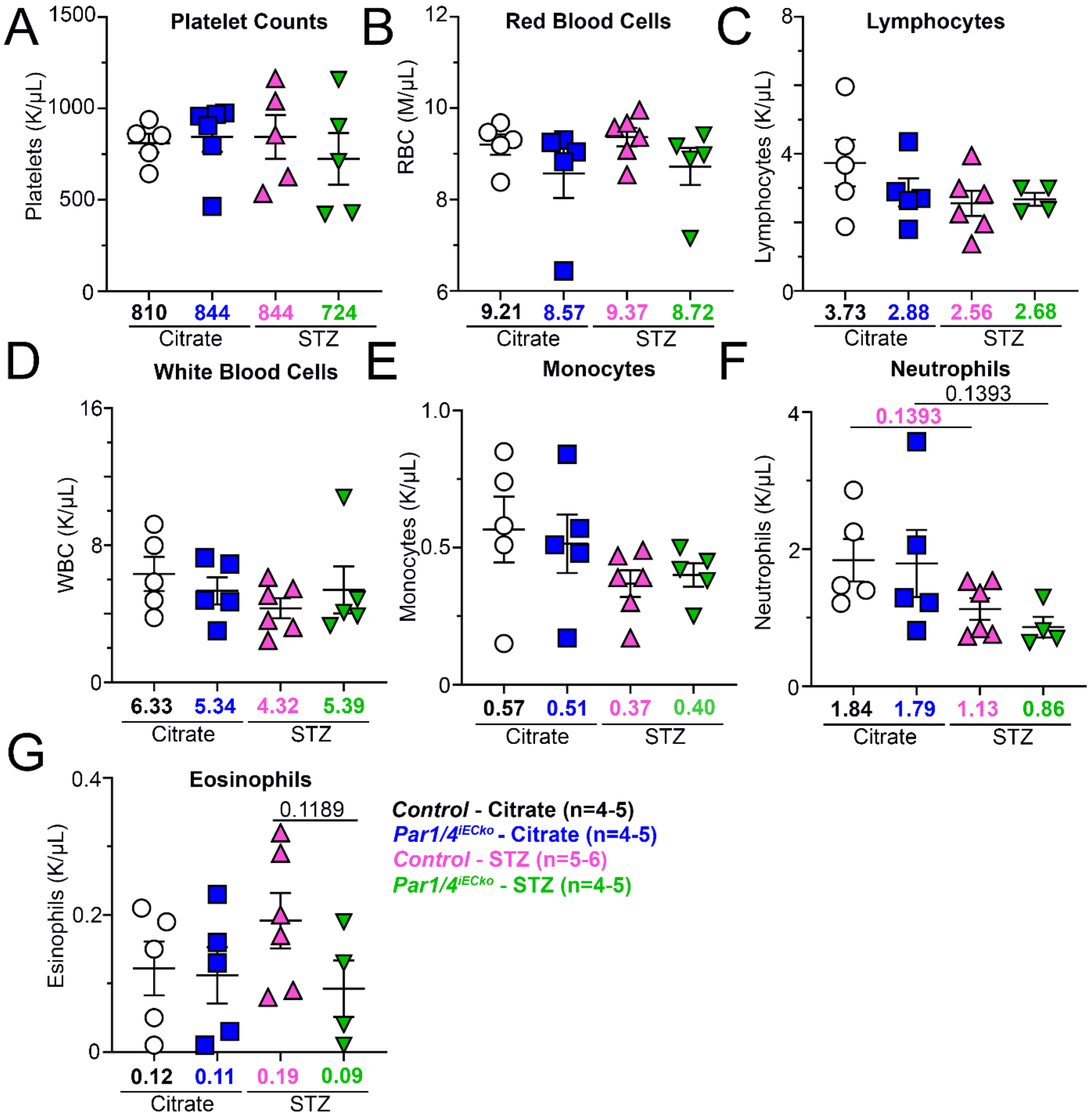

**<u>Figure S4.</u> Circulating blood counts of citrate- and STZ-treated *Par1/4*$^{iECko}$ mice and littermate controls.** Blood counts of (**A**) platelets, (**B**) red blood cells, (**C**) lymphocytes, (**D**) white blood cells, (**E**) monocytes, (**F**) neutrophils, and (**G**) eosinophils. <u>Statistics</u>: (F) Analyzed by Welch ANOVA followed by a 2-stage linear step-up procedure of Benjamini, Krieger, and Yekutieli for multiple comparison testing; q values are displayed in the graphs. (G) Analyzed by two-way ANOVA followed by a 2-stage linear step-up procedure of Benjamini, Krieger, and Yekutieli for multiple comparison testing; q values are displayed in the graphs. All scatter dot plots show mean ± SEM. (n=4-6 mice per group)

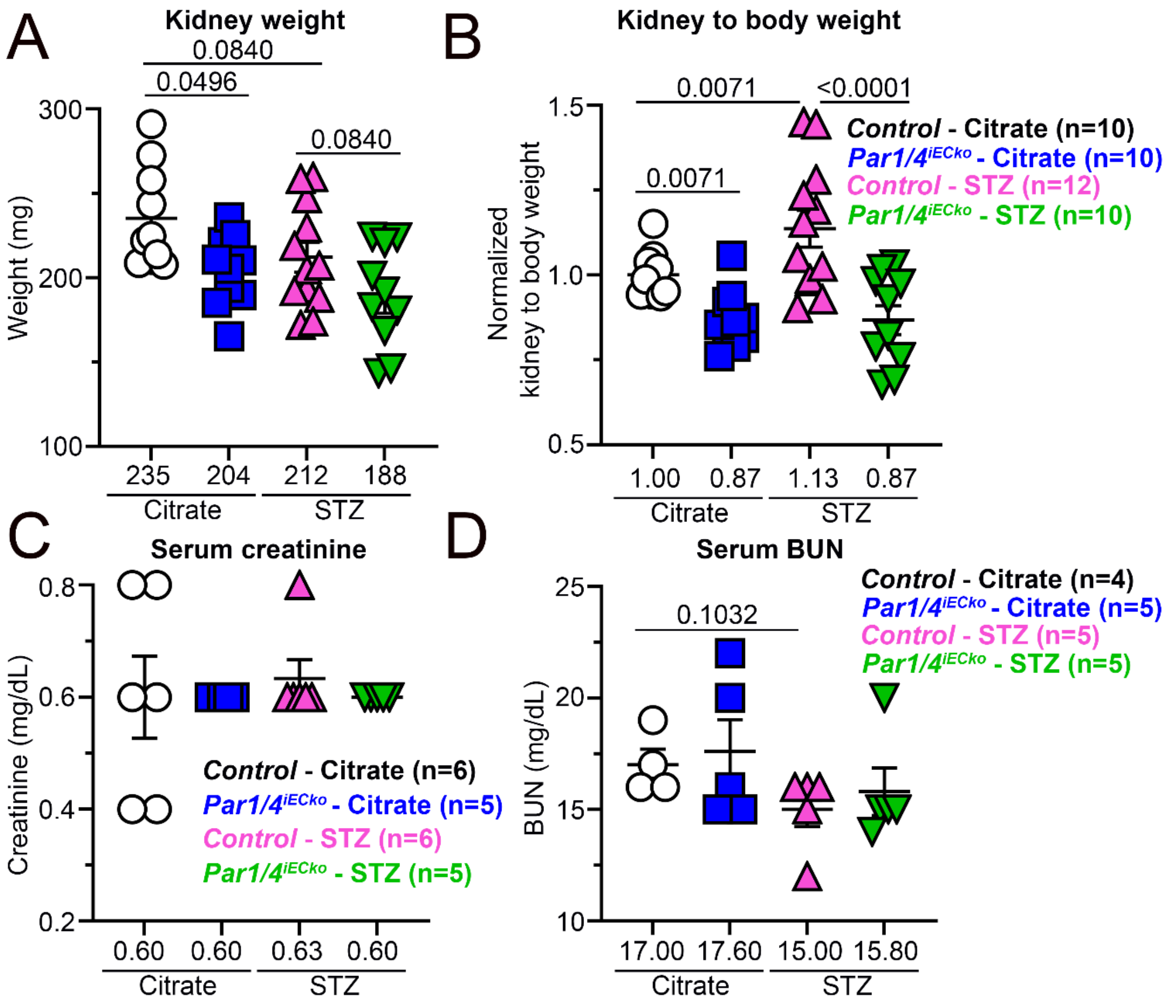

**<u>Figure S5.</u> STZ-treated *Par1/4*$^{iECko}$ mice display a reduction in some pathologies of renal dysfunction.** **(A)** Kidney weights and (**B**) kidney-to-body weights were assessed at the end of the study (12 weeks following STZ treatment) from citrate- and STZ-treated *Par1/4*$^{iECko}$ mice and littermate controls. (**C**) creatinine and (**D**) blood urea nitrogen (BUN) levels were assessed from serum. <u>Statistics</u>: (A-B) Analyzed by two-way ANOVA followed by a 2-stage linear step-up procedure of Benjamini, Krieger, and Yekutieli for multiple comparison testing; q values are displayed in the graphs. (n=10-12). (D) Analyzed by Welch ANOVA followed by a 2-stage linear step-up procedure of Benjamini, Krieger, and Yekutieli for multiple comparison testing; q values are displayed in the graphs. (n=4-5). All scatter dot plots show mean ± SEM.

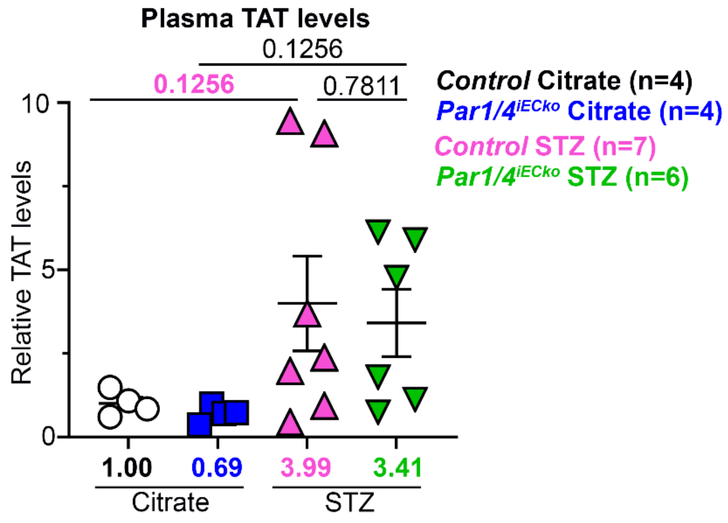

**<u>Figure S6.</u> Thrombin levels are modestly increased in mice at 12 weeks after STZ induction.** Plasma TAT (thrombin-antithrombin complex) was measured in mice at the end of the study. All values were normalized to citrate-treated control mice. Analyzed by Welch ANOVA followed by a 2-stage linear step-up procedure of Benjamini, Krieger, and Yekutieli for multiple comparison testing; q values are displayed in the graphs. All scatter dot plots show mean ± SEM. (n=4-7 mice per group)

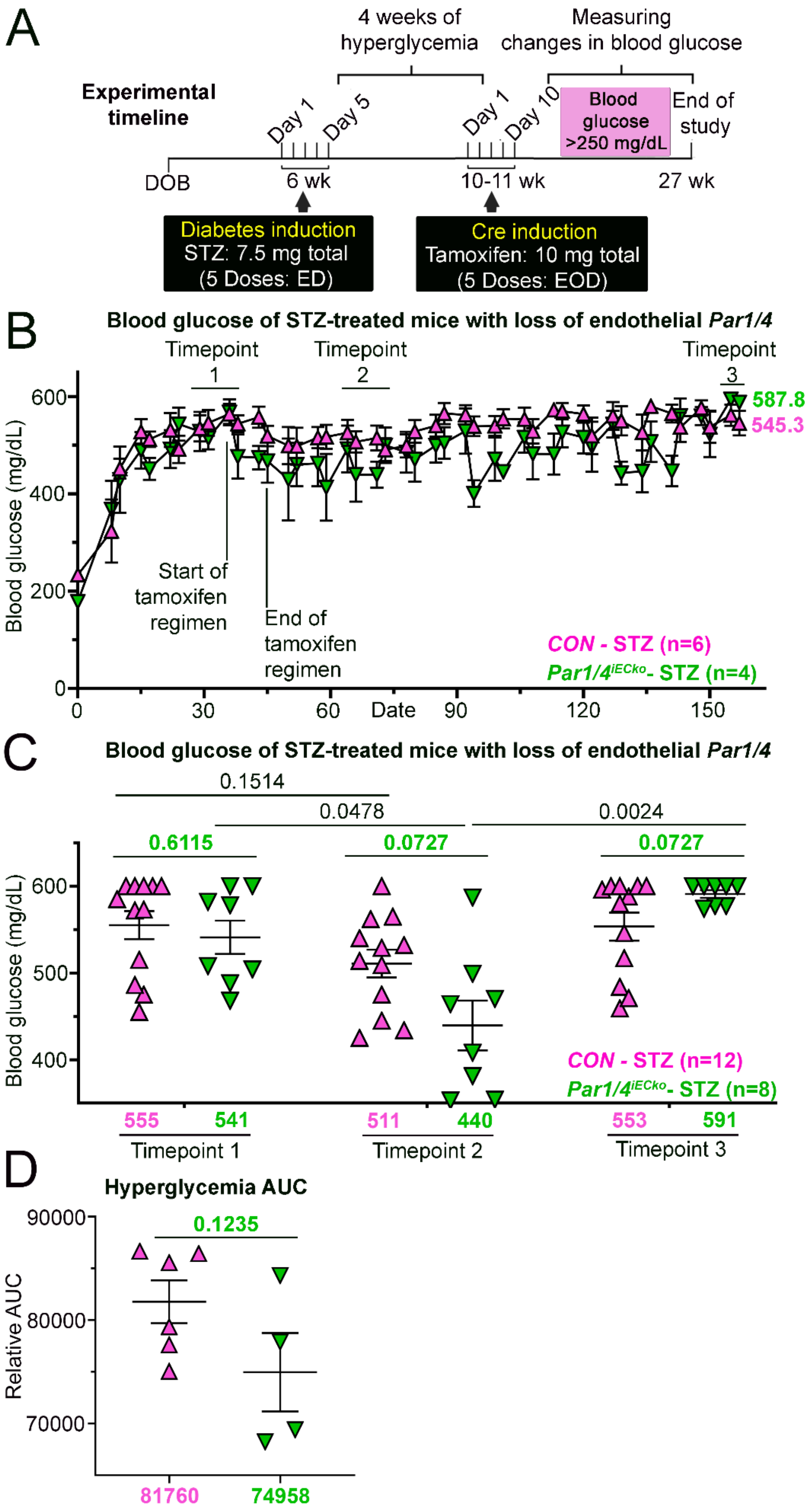
A
Experimental timeline
4 weeks of hyperglycemia
Measuring changes in blood glucose
Day 1
Day 5
Day 1
Day 10
Blood glucose >250 mg/dL
End of study
DOB
6 wk
10-11 wk
27 wk
Diabetes induction
STZ: 7.5 mg total
(5 Doses: ED)
Cre induction
Tamoxifen: 10 mg total
(5 Doses: EOD)
B
Blood glucose of STZ-treated mice with loss of endothelial Par1/4
Timepoint 1
Timepoint 2
Timepoint 3
587.8
545.3
Start of tamoxifen regimen
End of tamoxifen regimen
CON - STZ (n=6)
Par1/4iECko- STZ (n=4)
Blood glucose (mg/dL)
Date
C
Blood glucose of STZ-treated mice with loss of endothelial Par1/4
0.1514
0.0478
0.0024
0.6115
0.0727
0.0727
CON - STZ (n=12)
Par1/4iECko- STZ (n=8)
Blood glucose (mg/dL)
555
541
Timepoint 1
511
440
Timepoint 2
553
591
Timepoint 3
D
Hyperglycemia AUC
0.1235
Relative AUC
81760
74958

**<u>Figure S7</u>. Loss of endothelial PAR1 and PAR4 transiently reduces hyperglycemia in diabetic mice.** **(A)** Outline of experimental procedures performed on mice. DOB: date of birth; EOD: every other day; ED: every day. (n=4-6) **(B)** Blood glucose levels of STZ-treated mice prior to and following tamoxifen treatment to induce deletion of *Par1* and *Par4* in ECs. (**C**) Blood glucose is compared from three different time points: (Timepoint 1) before tamoxifen-induced deletion of *Par1* and *Par4*, (Timepoint 2) shortly after gene deletion, and (Timepoint 3) at the end of the study. **(D)** Area under the curve (AUC) for STZ-treated $Par1/4^{iECko}$ and littermate controls. <u>Statistics</u>: (C) Comparisons between control and $Par1/4^{iECko}$ were analyzed by Welch ANOVA followed by a 2-stage linear step-up procedure of Benjamini, Krieger, and Yekutieli for multiple comparison testing. Comparisons between $Par1/4^{iECko}$ at multiple different time points were analyzed by a Friedman test followed by a 2-stage linear step-up procedure of Benjamini, Krieger, and Yekutieli for multiple comparison testing; q values are displayed in the graphs. (n=8-12). A Wilcoxon matched-pair signed-rank test analyzed comparisons between control mice at two different time points followed by a 2-stage linear step-up procedure of Benjamini, Krieger, and Yekutieli for multiple comparison testing; q values are displayed in the graphs. (n=8-12). (D) Analyzed by unpaired t-test; p values are displayed in the graphs. (n=4-6). All scatter dot plots show mean ± SEM. CON: control.

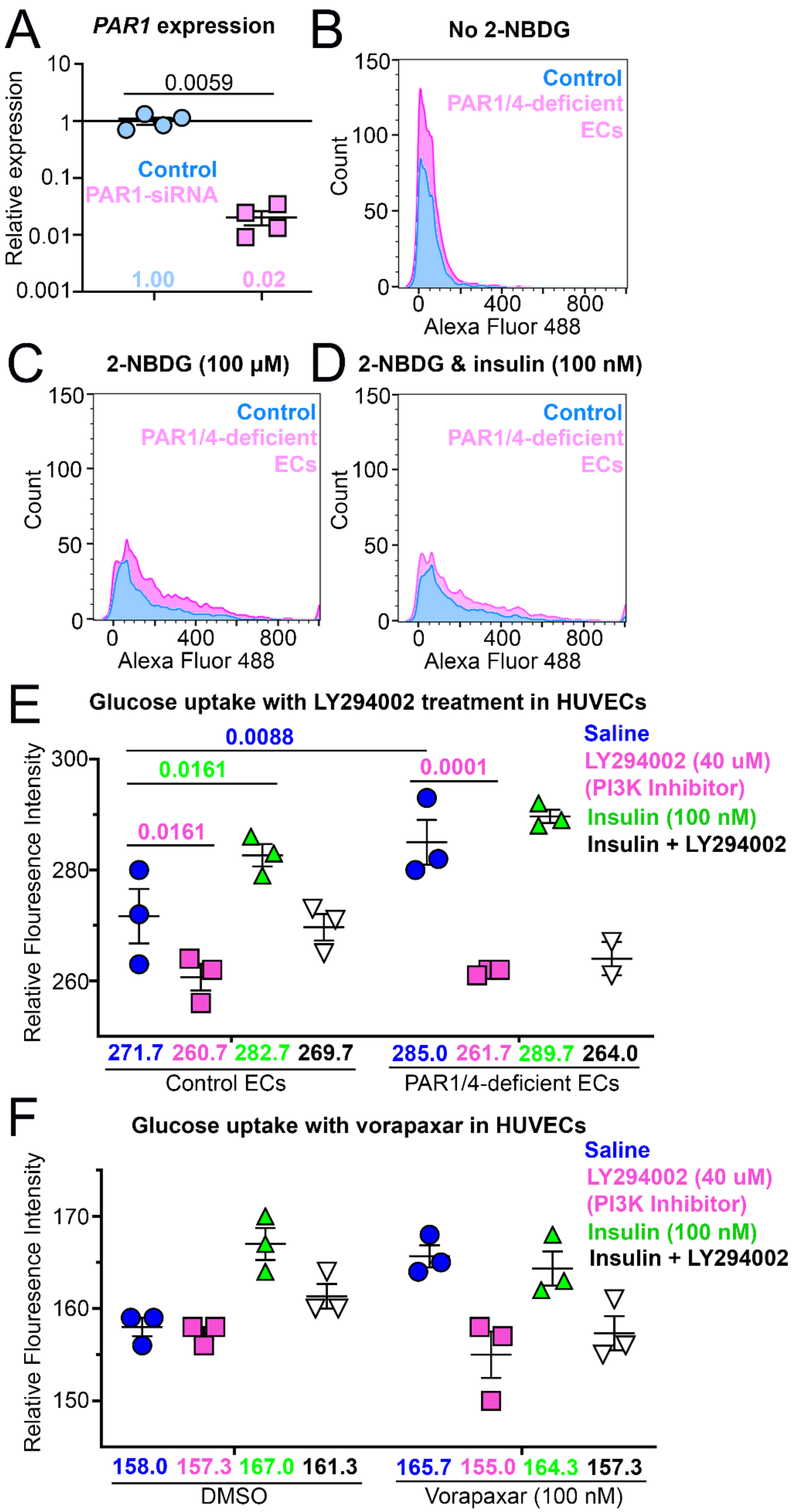
A
PAR1 expression
Relative expression
0.0059
Control
PAR1-siRNA
1.00
0.02
B
No 2-NBDG
Count
Control
PAR1/4-deficient ECs
Alexa Fluor 488
C
2-NBDG (100 µM)
Count
Control
PAR1/4-deficient ECs
Alexa Fluor 488
D
2-NBDG & insulin (100 nM)
Count
Control
PAR1/4-deficient ECs
Alexa Fluor 488
E
Glucose uptake with LY294002 treatment in HUVECs
Relative Flouresence Intensity
0.0088
0.0161
0.0161
0.0001
Saline
LY294002 (40 uM) (PI3K Inhibitor)
Insulin (100 nM)
Insulin + LY294002
271.7 260.7 282.7 269.7
Control ECs
285.0 261.7 289.7 264.0
PAR1/4-deficient ECs
F
Glucose uptake with vorapaxar in HUVECs
Relative Flouresence Intensity
Saline
LY294002 (40 uM) (PI3K Inhibitor)
Insulin (100 nM)
Insulin + LY294002
158.0 157.3 167.0 161.3
DMSO
165.7 155.0 164.3 157.3
Vorapaxar (100 nM)

541

**Figure S8.** **PAR1/4-deficient ECs have increased PI3K-mediated glucose uptake. (A)** *PAR1* expression at 48 hours following PAR1-siRNA treatment. Graphs of fluorescence intensity of control and PAR1/4-deficient ECs with (**B**) no treatment of 2-NBDG (a glucose analog), (**C**) treatment of 2-NBDG, and (**D**) treatment of 2-NBDG with insulin. **(E)** Flow cytometry analysis for uptake of 2-NBDG in control and PAR1/4-deficient HUVECs treated with saline (blue), PI3K inhibitor (LY294002; pink), insulin (green), and insulin plus LY294002 (black). (**F**) Flow cytometry analysis for 2-NBDG uptake in control and vorapaxar (PAR1 inhibitor)-treated HUVECs treated with saline (blue), LY294002 (pink), insulin (green), and insulin plus LY294002 (black). Statistics: (E) Analyzed by a Welch t-test; p values are displayed in the graphs. (**F**) Analyzed by 3-way ANOVA followed by a 2-stage linear step-up procedure of Benjamini, Krieger, and Yekutieli for multiple comparison testing; q values are displayed in the graphs. (n=3 technical replicates). All scatter dot plots show mean ± SEM.

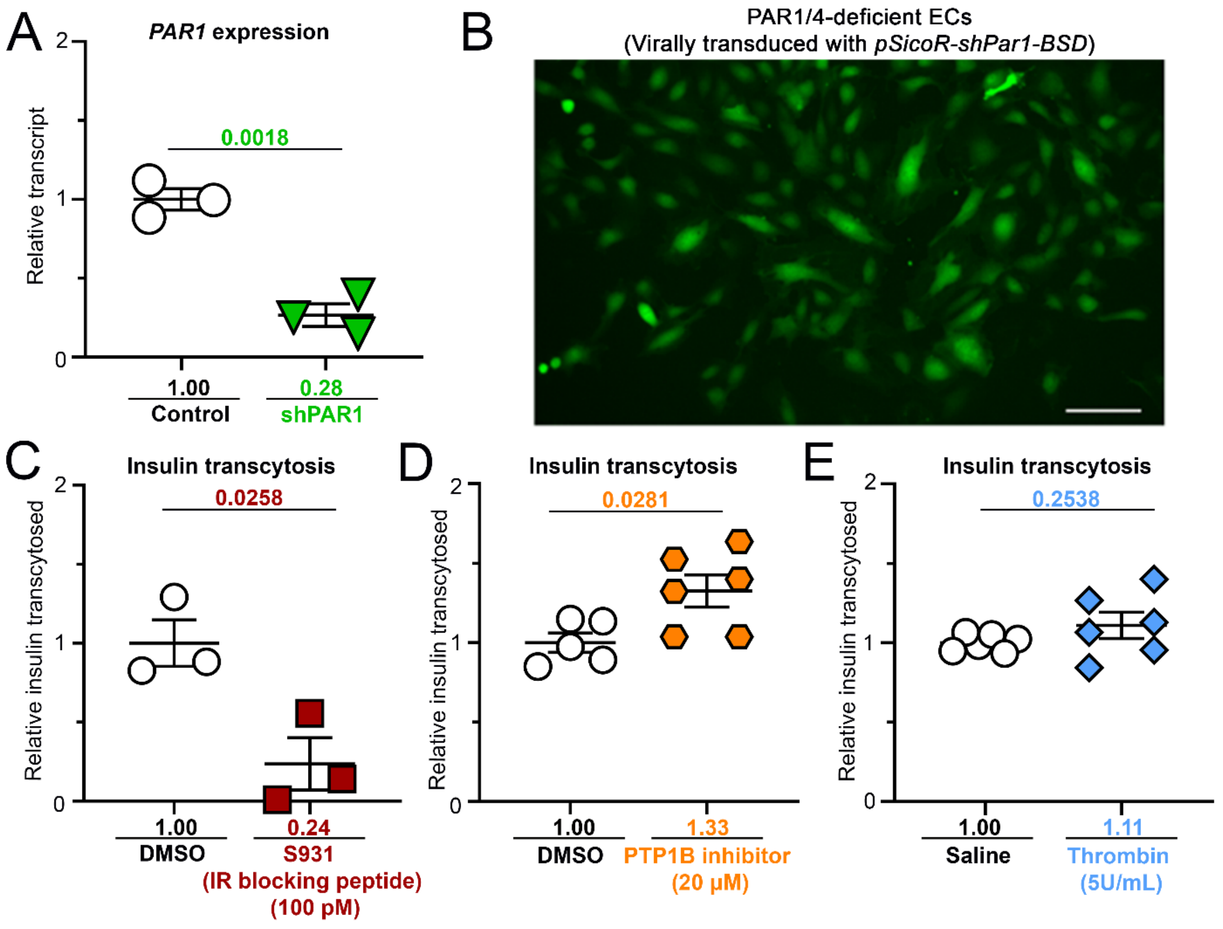

**<u>Figure S9.</u> Insulin transcytosis by ECs. (A)** *PAR1* expression following a PAR1-shRNA treatment. **(B)** GFP fluorescence of blasticidin (BSD) selected HUVEC colonies treated with PAR1-shRNA lentivirus; viruses carry eGFP-P2A-BSD cassettes resulting in eGFP mediated fluorescence. **(C)** An IR-blocking peptide (S931) reduced insulin transcytosis. (n=3 replicates per group). (**D**) PTP1B inhibition (JTT-551) increased insulin transcytosis. (n=5-6 replicates per group). (**E**) Insulin transcytosis of saline-treated control and thrombin-treated (5U/mL) ECs. (n=6 replicates per group). <u>Statistics</u>: (A-D) Analyzed by an unpaired t-test; p values are displayed in the graphs. (E) Analyzed by Welch t-test; p values are displayed in the graphs. All scatter dot plots show mean ± SEM.

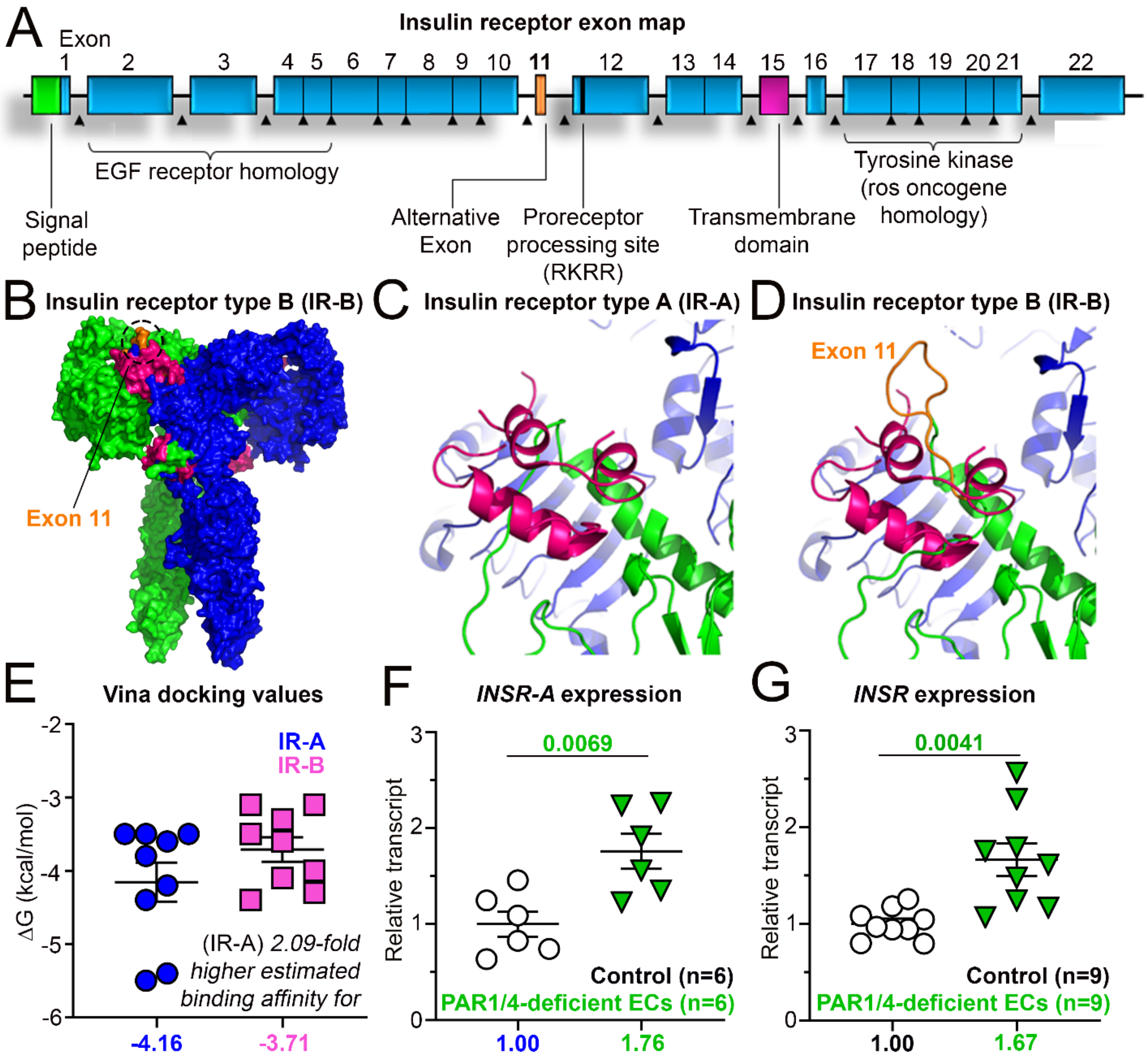

**<u>Figure S10.</u> Dual loss of PAR1 and PAR4 in HUVECs increases the expression of the *INSR-A* spliceform.** (**A**) Exon map of the *INSR* displaying alternative exon 11. (**B**) *In-silico* model of IR type B (IR-B), which contains residues of exon 11 (orange) that are shown to be present in two of the four insulin binding sites. Each monomer of the insulin receptor is shown in blue and green. Four insulins (pink) are shown bound to the insulin receptor. *In-silico* model was generated using a backbone from Uchikawa and colleagues (14). (**C**) Magnified view of the insulin binding site for IR-A and (**D**) IR-B. (**E**) Gibbs free energies of insulin binding to the insulin binding sites are seen in (C) and (D) for IR-A and IR-B, respectively. Data were generated using AutoDock Vina. Using the equation $\Delta G = -RTln(K_{eq})$, we can calculate the binding affinities of insulin to IR-A and IR-B, respectively. (**F**) *INSR-A* and (**G**) *INSR* expression in control and PAR1/4-deficient HUVECs. (n=6-9). Statistics: (F) Analyzed by unpaired t-test; p values are displayed in the graphs. (G) Analyzed by Welch t-test; p values are displayed in the graphs. All scatter dot plots show mean ± SEM.

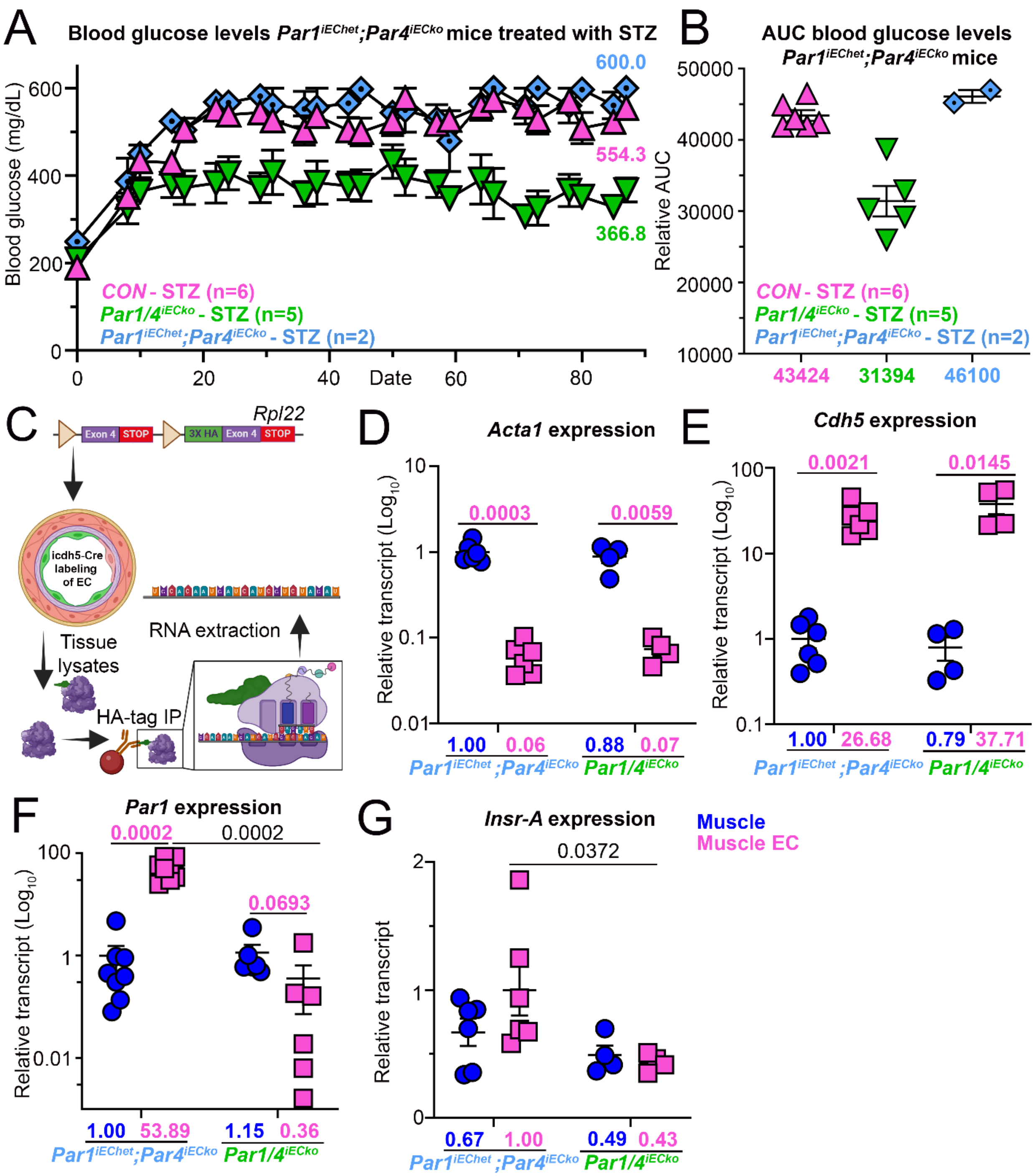
A
Blood glucose levels Par1iEChet;Par4iECko mice treated with STZ
Blood glucose (mg/dL)
600.0
554.3
366.8
CON - STZ (n=6)
Par1/4iECko - STZ (n=5)
Par1iEChet;Par4iECko - STZ (n=2)
Date
B
AUC blood glucose levels
Par1iEChet;Par4iECko mice
Relative AUC
CON - STZ (n=6)
Par1/4iECko - STZ (n=5)
Par1iEChet;Par4iECko - STZ (n=2)
43424
31394
46100
C
Rpl22
Exon 4
STOP
3X HA
Exon 4
STOP
icdh5-Cre labeling of EC
Tissue lysates
RNA extraction
HA-tag IP
D
Acta1 expression
Relative transcript (Log10)
0.0003
0.0059
1.00
0.06
0.88
0.07
Par1iEChet;Par4iECko
Par1/4iECko
E
Cdh5 expression
Relative transcript (Log10)
0.0021
0.0145
1.00
26.68
0.79
37.71
Par1iEChet;Par4iECko
Par1/4iECko
F
Par1 expression
Relative transcript (Log10)
0.0002
0.0002
0.0693
1.00
53.89
1.15
0.36
Par1iEChet;Par4iECko
Par1/4iECko
G
Insr-A expression
Relative transcript
0.0372
0.67
1.00
0.49
0.43
Par1iEChet;Par4iECko
Par1/4iECko
Muscle
Muscle EC

**<u>Figure S12.</u> *$Par1/4^{iECko}$* mice display decreased expression of the *Insr-A* spliceform in muscle endothelium. (A)** Blood glucose levels of STZ-treated male *$Par1/4^{iECko}$* and *$Par1^{iEChet}$;$Par4^{iECko}$* (control) mice. **(B)** Area under the curve (AUC) values for hyperglycemia of mice in panel A. (**C**) Schematic of endothelial TRAP isolation. **(D-G)** Samples were collected from the quadricep muscles of *$Par1/4^{iECko}$* and *$Par1^{iEChet}$;$Par4^{iECko}$* mice 4 weeks after tamoxifen treatment for gene deletion. (D) *Acta1* (E) *Cdh5* (F) *Par1* (G) *Insr-A* expression compared between muscle and muscle endothelium of *$Par1/4^{iECko}$* and *$Par1^{iEChet}$;$Par4^{iECko}$* mice. In (D-F) expression was normalized to the muscle samples of *$Par1^{iEChet}$;$Par4^{iECko}$* mice and data are shown on a $Log_{10}$ scale. In (G) expression was normalized to the muscle endothelium samples of *$Par1^{iEChet}$;$Par4^{iECko}$* mice. <u>Statistics</u>: All panels were analyzed by Welch ANOVA followed by a 2-stage linear step-up procedure of Benjamini, Krieger, and Yekutieli for multiple comparison testing; q values are displayed in the graphs. All scatter dot plots show mean ± SEM. (n=4-8 mice)

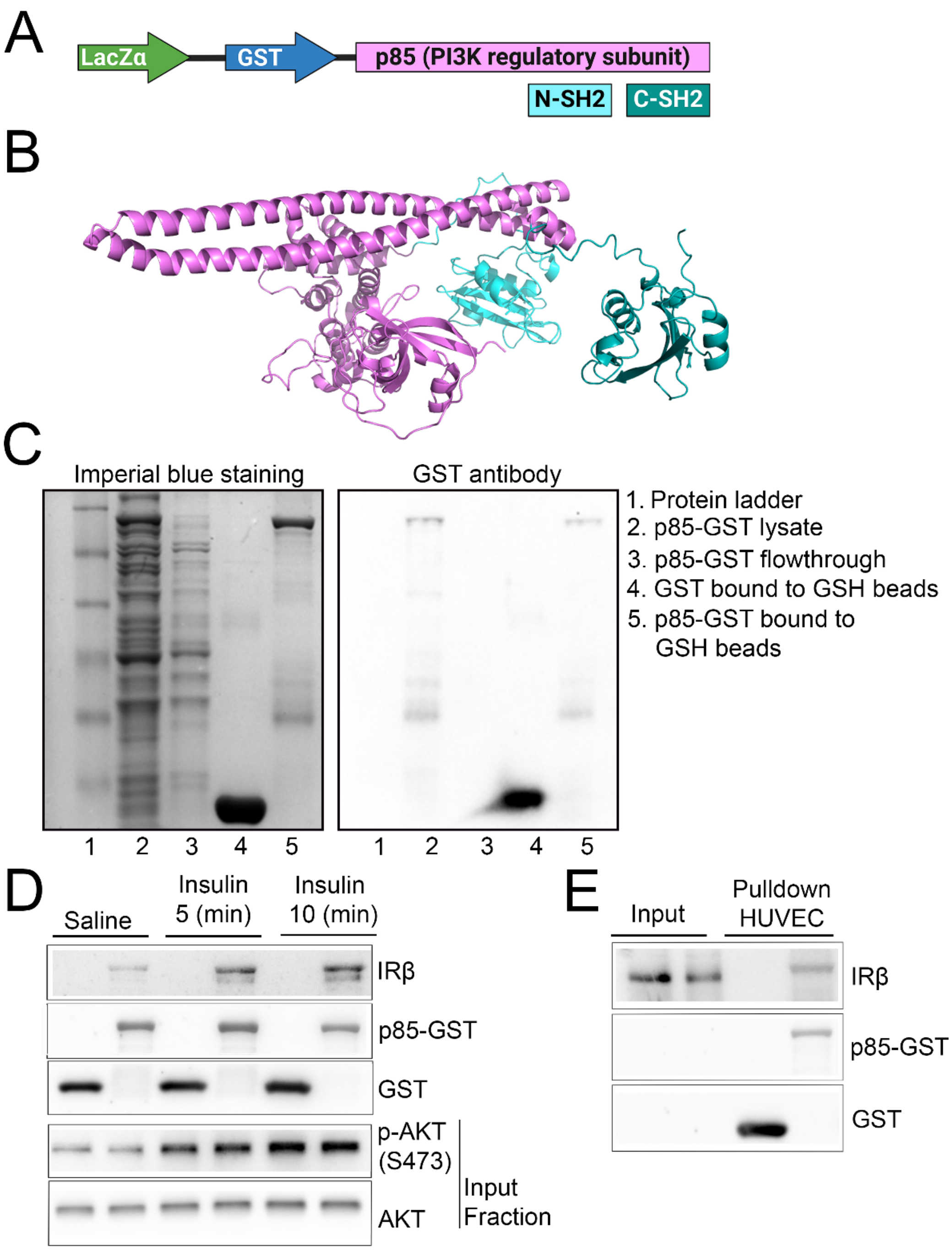

**<u>Figure S12.</u> Validation of the p85-GST probe. (A)** Schematic of the p85-GST probe. (**B**) AlphaFold model of p85 with N-SH2 and C-SH2 domains colored. (**C**) Imperial blue staining and a GST immunoblot of GST and GST-p85 probe isolated from BL21 cells. (**D**) p85-GST pulldowns of quadriceps lysates (500 µg) from overnight-fasted mice treated intraperitoneally with saline and insulin (5U/kg) for 5 and 10 minutes, probed with IRβ. IR phosphorylation is increased in insulin-treated samples. The two bottom panels in (D) are muscle lysates of samples in the upper panel, showing increased p-AKT (Ser 473) in response to insulin treatment that verifies IR activation. (**E**) p85-GST pulldowns of HUVEC lysate (100 μg) showing basal IR phosphorylation of ECs.

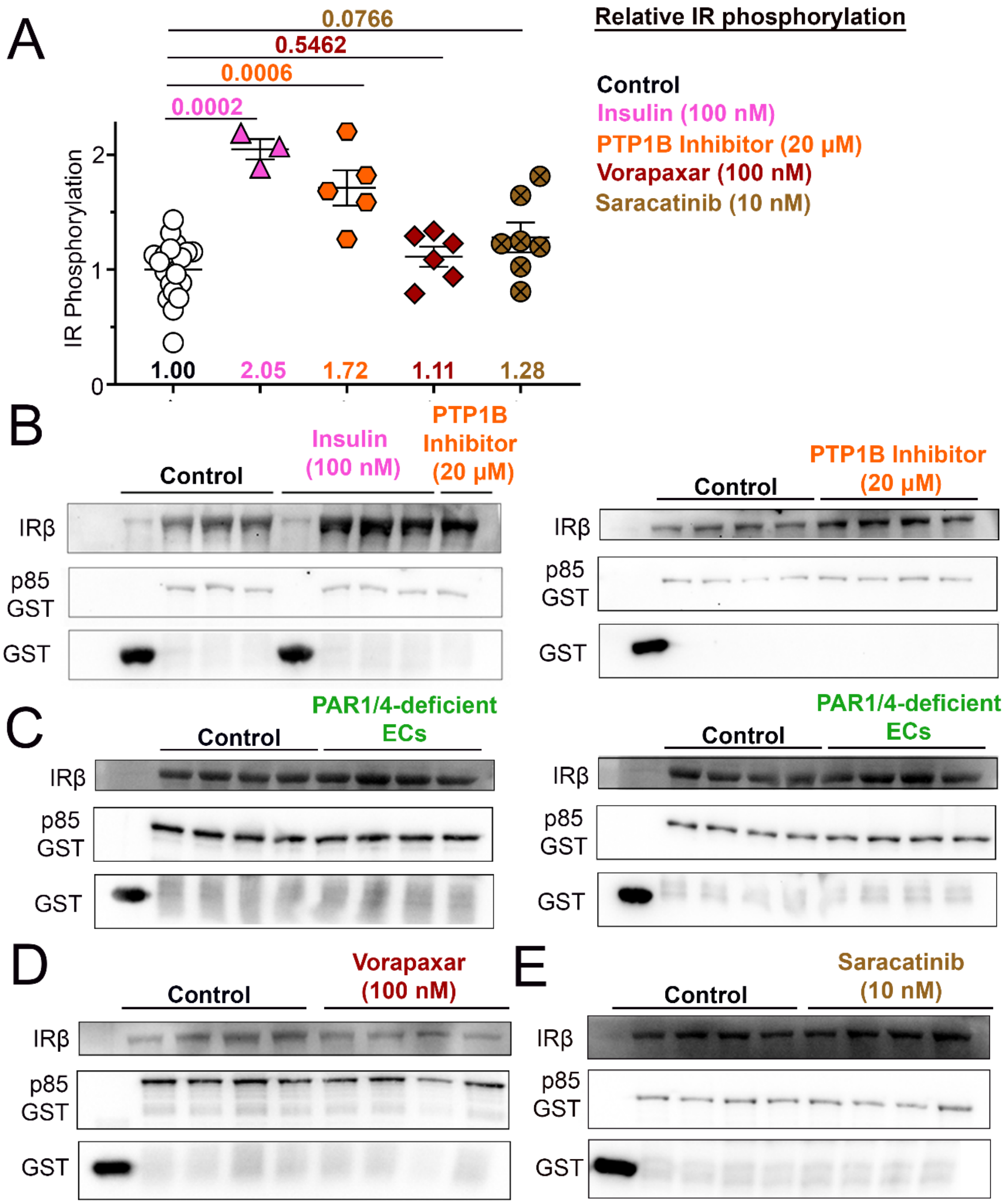

<u>**Figure S13.**</u> **Measurements of IR activity with p85-GST pulldown assays in treated HUVECs.** (A) Quantification of relative IR activity as measured by p85-GST pulldowns of HUVECs treated with insulin, a PTP1B inhibitor (JTT-551), a PAR1 inhibitor (vorapaxar; 100 nM), or a c-Src inhibitor (saracatinib; 10 nM). (**B**) Immunoblots of p85-GST pulldowns of HUVECs treated with insulin and JTT-

551. Immunoblots of p85-GST pulldowns of (**C**) control and PAR1/4-deficient ECs, (**D**) control and vorapaxar-treated ECs, and (**E**) control and saracatinib-treated ECs. Statistics: (A) Insulin- and PTP1B inhibitor-treated samples were analyzed by one-way ANOVA; q values are displayed in the graphs. (n=3-7 replicates per group). Vorapaxar- and saracatinib-treated samples were analyzed using an unpaired t-test; p values are displayed in the graphs. (n=6-7 technical replicates per group). All scatter dot plots show mean ± SEM.

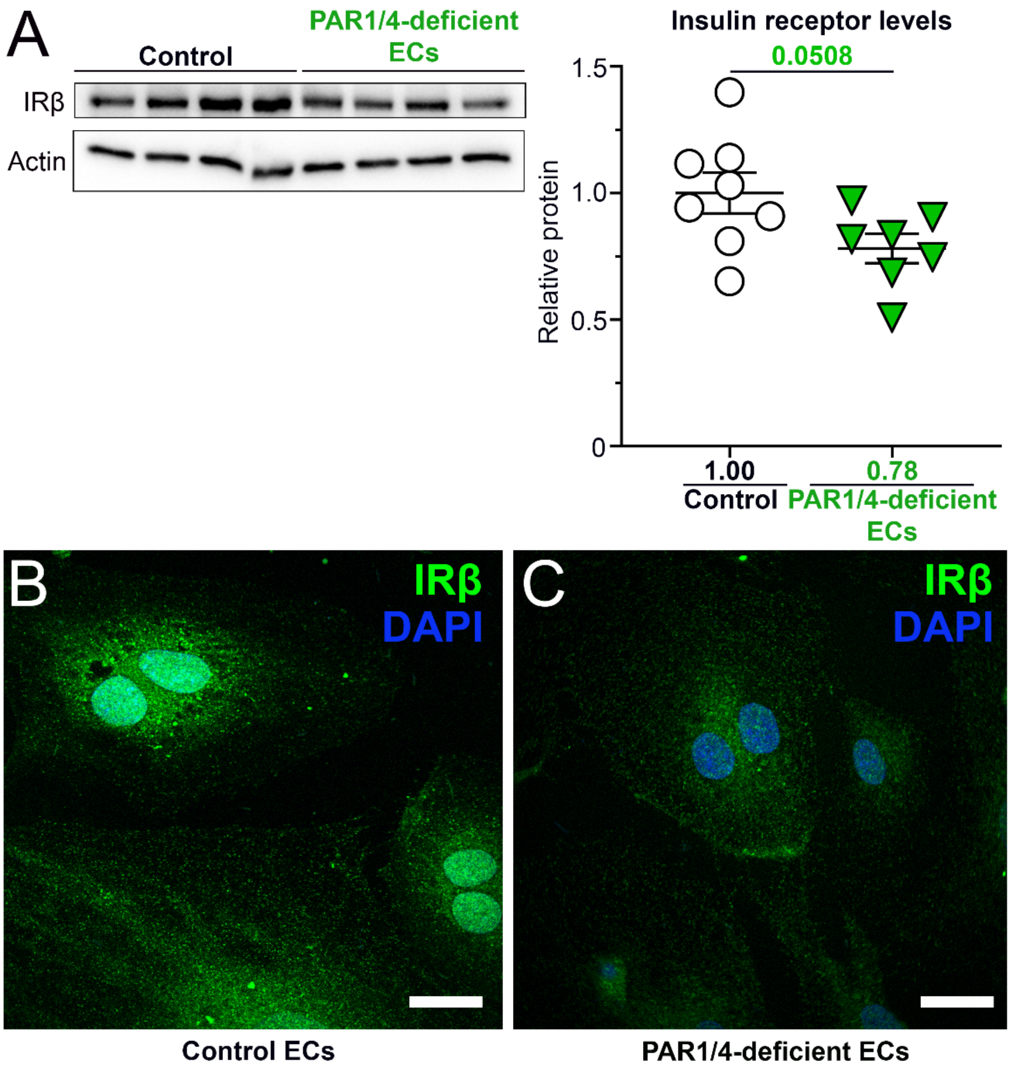

**<u>Figure S14.</u> IR is moderately reduced in PAR1/4-deficient ECs. (A)** Comparison of IR protein levels between control and PAR1/4-deficient HUVECs by immunoblot. **(B, C)** Representative confocal immunofluorescence images depicting insulin receptor expression in (B) control and (C) PAR1/4-deficient HUVECs. Scale bar: 20 µm. <u>Statistics</u>: (A) Samples are normalized to β-actin. (n=7-8 technical replicates). Data were analyzed using an unpaired t-test; p values are displayed in the graphs. All scatter dot plots show mean ± SEM.

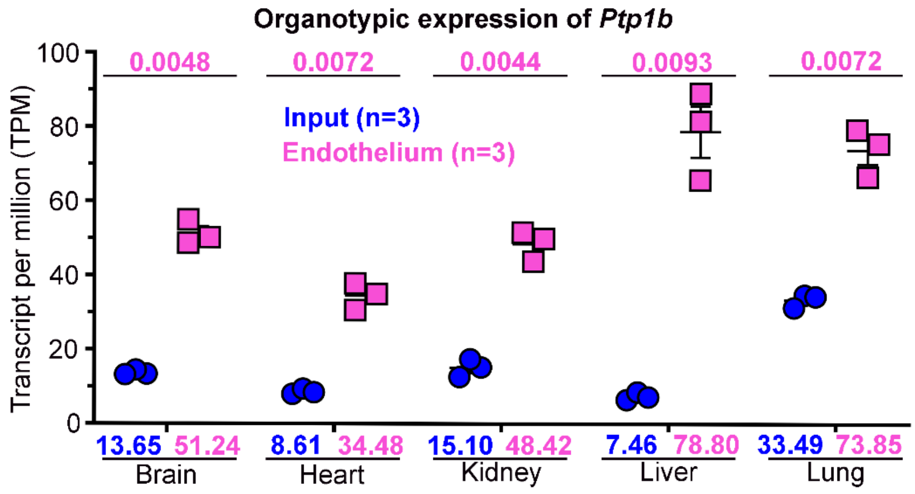

**<u>Figure S15.</u> PTP1B is enriched in ECs.** An analysis of TRAP data generated by Cleuren et al (21) reveals *Ptp1b* expression in input lysates and ECs of different murine tissue beds. <u>Statistics</u>: Data was analyzed using Welch's t-test followed by a 2-stage linear step-up procedure of Benjamini, Krieger, and Yekutieli for multiple comparison testing; q values are displayed in the graphs. All scatter dot plots show mean ± SEM. (n=3)

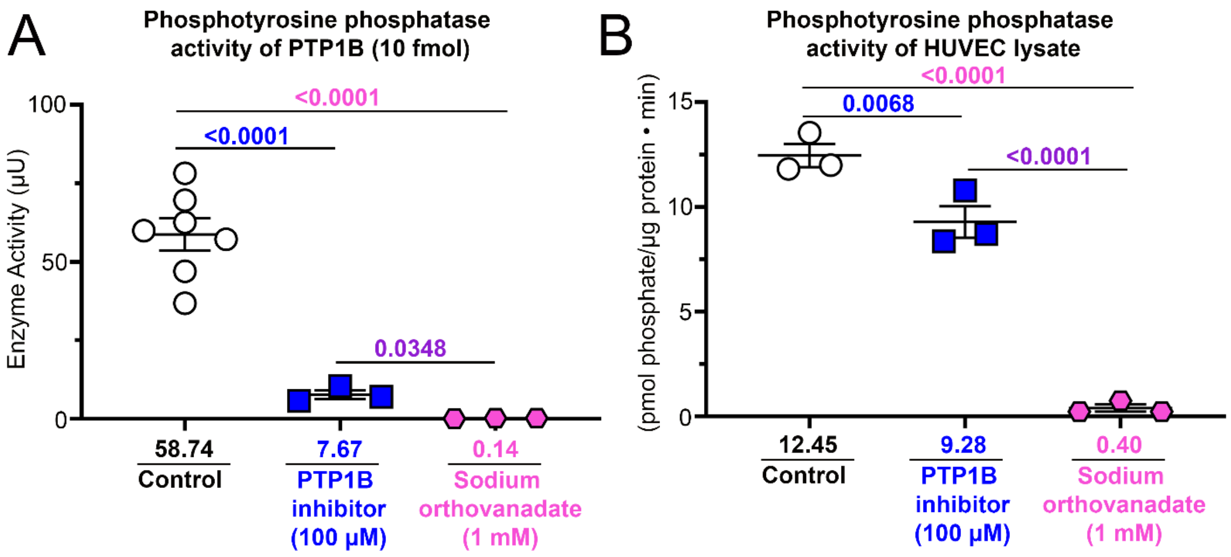

**<u>Figure S16.</u> Phosphotyrosine phosphatase activity of PTP1B and EC lysates with various phosphatase inhibition.** Comparison of the enzymatic activity of (**A**) 10 femtomoles of PTP1B or (**B**) HUVEC lysates treated with an allosteric inhibitor of PTP1B and a broad-spectrum phosphatase inhibitor (sodium orthovanadate). <u>Statistics</u>: (A) analyzed using Welch's ANOVA followed by a 2-stage linear step-up procedure of Benjamini, Krieger, and Yekutieli for multiple comparison testing; q values are displayed in the graphs. (B) analyzed using a one-way ANOVA followed by a 2-stage linear step-up procedure of Benjamini, Krieger, and Yekutieli for multiple comparison testing; q values are displayed in the graphs. All scatter dot plots show mean ± SEM. (n=3-7)

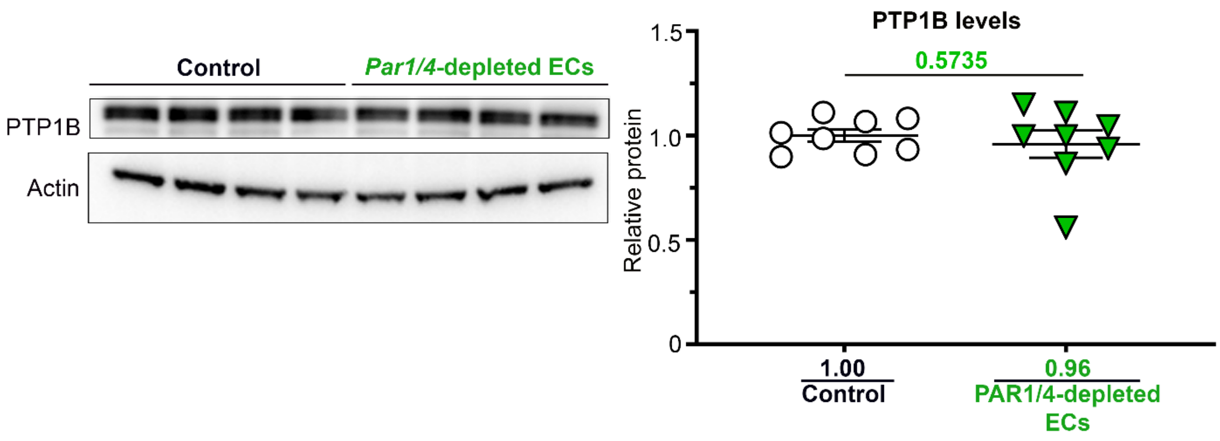

**<u>Figure S17.</u>** **PTP1B levels are unchanged in PAR1/4-deficient ECs.** Comparison of PTP1B protein levels between control and PAR1/4-deficient HUVECs by immunoblot. <u>Statistics</u>: Samples are normalized to β-actin. Data were analyzed using a Welch t-test; p values are displayed in the graphs. All scatter dot plots show mean ± SEM. (n=8)

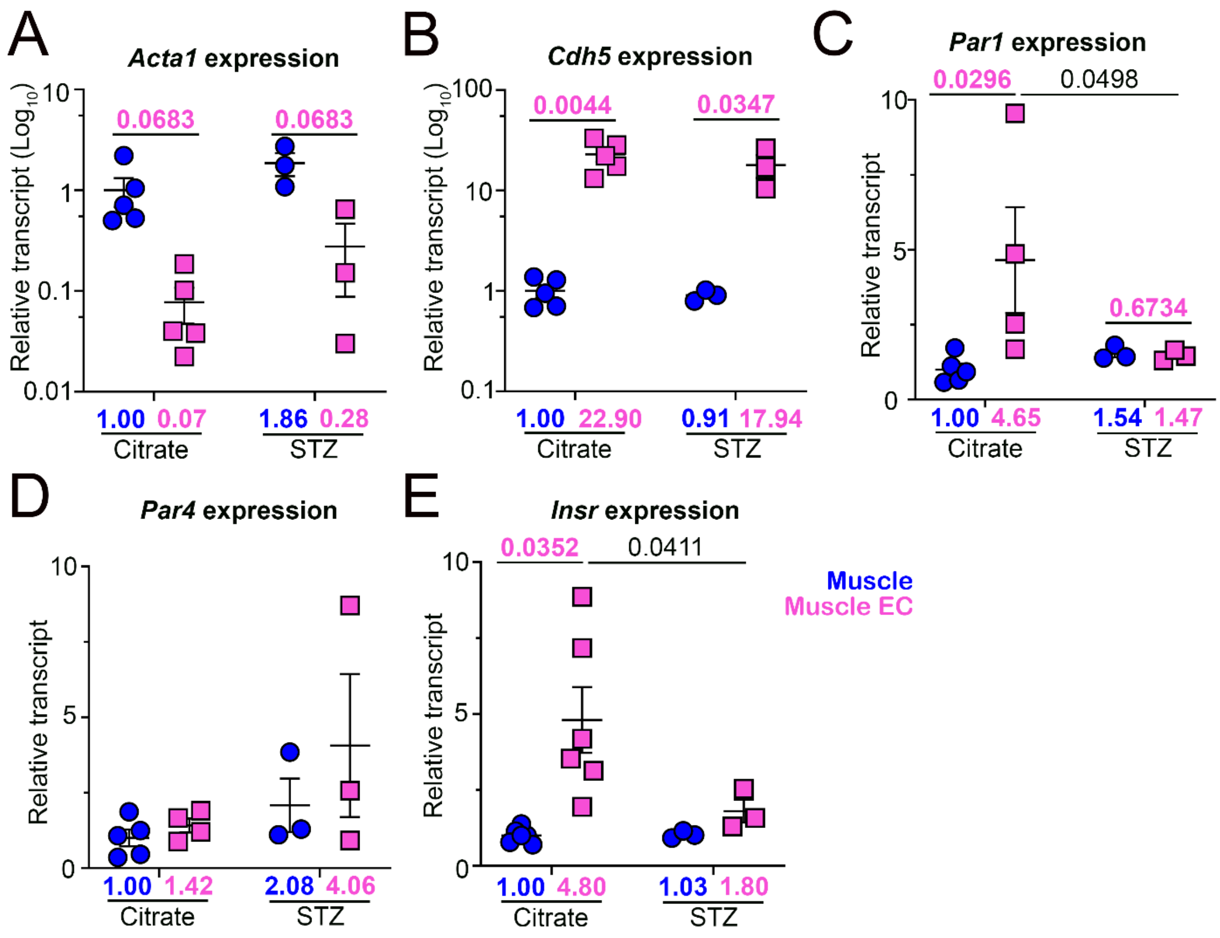

**<u>Figure S18.</u> Diabetic mice show reduced expression of *Insr* in muscle ECs.** The quadriceps of endothelial TRAP mice were collected following 16 weeks of hyperglycemia. **(A)** *Acta1* **(B)** *Cdh5* **(C)** *Par1* **(D)** *Par4* **(E)** *Insr* expression compared between muscle and the muscle endothelium of control and diabetic mice. Expression was normalized to the muscle samples of control mice. (n=3-6 mice per group). <u>Statistics</u>: (A, B, E) Analyzed by Welch ANOVA followed by a 2-stage linear step-up procedure of Benjamini, Krieger, and Yekutieli for multiple comparison testing; q values are displayed in the graphs. (C) Analyzed by two-way ANOVA followed by a 2-stage linear step-up procedure of Benjamini, Krieger, and Yekutieli for multiple comparison testing; q values are displayed in the graphs. In (A-B) data are shown on a $Log_{10}$ scale. All scatter dot plots show mean ± SEM.

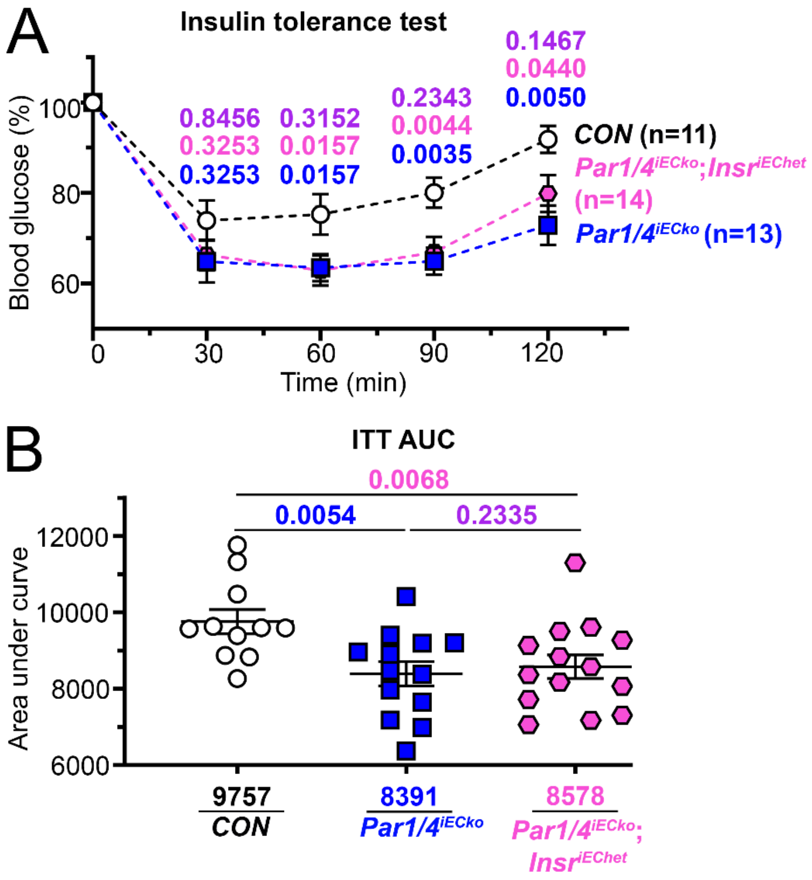

**<u>Figure S19.</u> Endothelial-specific deletion of one allele of *Insr* in *Par1/4*$^{iECko}$ mice subtly alters the loss of insulin sensitivity observed in *Par1/4*$^{iECko}$ mice.**

(**A**) Insulin tolerance test (ITT) normalized to baseline reading (insulin administered: 1 U/kg) between control (CON), *Par1/4*$^{iECko}$, and *Par1/4*$^{iECko}$;*Insr*$^{iEChet}$ mice. (**B**) The area under the curve (AUC) is calculated for each ITT measurement in A. <u>Statistics</u>: (A) Each time point was analyzed with a one-way ANOVA followed by a 2-stage linear step-up procedure of Benjamini, Krieger, and Yekutieli for multiple comparison testing; q values are displayed. (B) Analyzed with a one-way ANOVA followed by a 2-stage linear step-up procedure of Benjamini, Krieger, and Yekutieli for multiple comparison testing; q values are displayed in the graphs. n=11–14 mice per group. Blue text indicates a comparison between control and *Par1/4*$^{iECko}$ mice. Pink text indicates a comparison between control and *Par1/4*$^{iECko}$;*Insr*$^{iEChet}$ mice. Purple text indicates a comparison between *Par1/4*$^{iECko}$ and *Par1/4*$^{iECko}$;*Insr*$^{iEChet}$ mice.

**SUPPLEMENTAL TABLE 1 (ST1): Genotyping Primers**

| **Allele** | **Forward primer** | **Reverse primer** | **$T_m$ (C)** | **Bands (bp)** |
|---|---|---|---|---|
| *Par1* | 5’ - CATGGGGGAAGCTATCAGAA | 5’ - CCAAAACCGAGTCCAAAAGA | 55 | WT – 202<br>Flox – 236 |
| *Par4* | 5’ - GGGATGTTGTGGAGATTTGG | 5’ - ATGGCTTCCCCTGGTACTCT | 55 | WT – 431<br>Flox – 465 |
| *Insr* | 5’- GGGGCAGTGAGTATTTTGGA | 5’- GGGCCACTAAAGGATGTGTC | 55 | WT – 257<br>Flox – 297 |
| RiboTag | 5’ - GGGAGGCTTGCTGGATATG | 5’ - TTTCCAGACACAGGCTAAGT ACAC | 55 | WT – 243<br>Flox – 290 |
| *Cdh5(PAC) -Cre*$^{ERT2}$ | 5’ - TCCTGATGGTGCCTATCCTC | 5’ - CGAACCTGGTCGAAATCAGT | 55 | 473 |

**SUPPLEMENTAL TABLE 2 (ST2): qPCR primers**

**Mouse**

| Allele | Forward primer | Reverse primer | $T_m$ (C) | Amplicon Size (bp) |
|---|---|---|---|---|
| *Par1* | 5’-CTTCCCGCGTCCCTATGAG | 5’-CTGGTCAGATATCCGGAGGC | 60 | 247 |
| *Par4* | 5’-CCTGCCCTCTATGGGCTT | 5’-AGCAGAATGGTGGATGGCA | 60 | 104 |
| *Insr* | 5’-GCTGTGCCATTGCTGGTG | 5’-CAGCTCATGTAGCCTGGTCA | 60 | 120 |
| *Splice* | 5’-AGGAGCTGGAGGAGTCTTCA | 5’-CTACTGTCCTCGGCACCATT | 60 | 106 |
| *Acta1* | 5’-CCATCGGCAATGAGCGTTTC | 5’-CCCCCTGACATGACGTTGTT | 60 | 160 |
| *Cdh5* | 5’-AGAGTCCATCGCAGAGTC | 5’-CAGCCAGCATCTTGAACC | 60 | 107 |

**Human**

| Allele | Forward primer | Reverse primer | $T_m$ (C) | Amplicon Size (bp) |
|---|---|---|---|---|
| *PAR1* | 5’-CGCAGGCCAGAATCAAAAGC | 5’-CCGGTGTACACAGATGGGAC | 60 | 254 |
| *PAR4* | 5’-CACCGGAGGTGGTGATGACAG | 5’-ACATGTGACCATAGAGTGCGG | 60 | 389 |
| *INSR* | 5’-CTGTACCCCGGAGAGGTGT | 5’-GGGCCTCGTTTTGAACATC | 60 | 126 |
| *Splice* | 5’-CCTGAAGGAGCTGGAGGAGT | 5’-TAGGGTCCTCGGCACCAGT | 60 | 110 |

**SUPPLEMENTAL TABLE 3 (ST3): Major resources table**

**Animals (in vivo studies)**

| Species | Vendor or Source | Background Strain | Sex | Persistent ID / URL |
|---|---|---|---|---|
| Mouse | Generated for our lab by ViewSolid Biotech, Inc | $Par1^{flox}$ | M | |
| Mouse | Generated for our lab by ViewSolid Biotech, Inc | $Par4^{flox}$ | M | |
| Mouse | Jackson Laboratory | $Insr^{flox}$ | M | #006955 |
| Mouse | Jackson Laboratory | RiboTag | M | #029977 |
| Mouse | Taconic | $Cdh5(PAC)\text{-}Cre^{ERT2}$ | M | #13073 |

**Genetically Modified Animals (Not applicable)**

| | **Species** | **Vendor or Source** | **Background Strain** | **Other Information** | **Persistent ID / URL** |
|---|---|---|---|---|---|
| **Parent - Male** | | | | | |
| **Parent - Female** | | | | | |

**Primary Antibodies**

| Target antigen | Vendor or Source | Catalog # | Working concentration | Lot # | Persistent ID / URL |
|---|---|---|---|---|---|
| HA-Tag | CST | C29F4 | 0.825 ng/mL (IP); i.e., 1:80 | | RRID: AB_1549585 |
| Insulin | BioRad | 7F8 (E6E5) | 1 µg/mL, i.e. 1:1000 | | RRID: AB_1068697 |
| Insulin Receptor | CST | 23413S | 1:1000 (WB) 1:25 (ICC) | | RRID: AB_2924796 |
| Phospho-Akt Threonine 308 | CST | 9275S | 1:1000 | | RRID: AB_329828 |
| Phospho-Akt Serine 473 | CST | 9271S | 1:1000 | | RRID: AB_329825 |
| Akt | CST | 9272S | 1:1000 | | RRID: AB_329827 |
| GST | CST | 2622S | 4.4 ng/mL, i.e. 1:5000 | | RRID: AB_331670 |
| Actin | CST | 4967S | 16 ng/mL, i.e. 1:1000 | | RRID: AB_330288 |
| sheep anti-thrombin antibody | Affinity Biologicals | SAFII-AP | 2 ug/mL | | https://affinitybiologicals.com/products/prothrombin-factor-ii-polyclonal-antibody/ |

| | | | | | |
|---|---|---|---|---|---|
| sheep anti-human antithrombin | Affinity Biologicals | SAAT-APBIO | 0.5 ug/mL | | https://affinitybiologicals.com/products/atiii-antithrombin-polyclonal-antibody-affinity-purified-biotinylated/ |
| PTP1B | Proteintech | 11334-1-AP | 346 ng/mL | | RRID: AB_10642566 |

**Secondary Antibodies**

| Host and Target species | Fluorophore or conjugate | Vendor or Source | Catalog # | Working concentration | Lot # | Persistent ID / URL |
|---|---|---|---|---|---|---|
| Horse anti-mouse | HRP | CST | 7076S | 1:2000 | | RRID: AB_330924 |
| Goat anti-Rabbit | HRP | Sigma | A6667 | 1:2000 | | RRID: AB_258307 |
| Donkey anti-Rabbit | FITC | Thermo Fisher Scientific | 711-095-152 | 1 µg/mL, i.e. 1:500 | | AB_2535792 |

**DNA/cDNA Clones**

| Clone Name | Sequence (Provided in supplements) | Source / Repository | Persistent ID / URL |
|---|---|---|---|
| pSicoR | pSicoR.DNA | Addgene | https://www.addgene.org/11579/ |
| pSicoR-shPAR1 | pSicoR-shPAR1.DNA | Generated by authors | |
| pSicoR-BSD | pSicoR-BSD.DNA | Generated by authors | |
| pSicoR-shPAR1-BSD | pSicoR-shPAR1-BSD.DNA | Generated by authors | |
| GST-4T1 | GST-4T1.DNA | Gift from Raju Rajala, Ph.D. | |
| p85-GST-4T1 | p85-GST-4T1.DNA | Gift from Raju Rajala, Ph.D. | |
| psPAX2 | psPAX2.DNA | Addgene | https://www.addgene.org/12260/ |
| PMD2.G | PMD2.G.DNA | Addgene | https://www.addgene.org/12259/ |

**Cultured Cells**

| Name | Vendor or Source | Sex (F, M, or unknown) | Persistent ID / URL |
|---|---|---|---|
| Human Umbilical Vein Endothelial Cells (HUVEC) | ATCC, PCS-100-010 | unknown | https://www.atcc.org/products/pcs-100-010?matchtype=&network=x&device=c&adposition=&keyword=&gad_source=1&gclid=Cj0KCQjwrp-3BhDgARIsAEWJ6Szj6tdAr2GPx2x5YCiGwf0OScN4X5hlmcoozBqhdJjCt8okKfxKxE4aAh_wEALw_wcB |
| Human Embryonic Kidney 293 (HEK-293) | ATCC, CRL-1573 | unknown | https://www.atcc.org/products/crl-1573 |

**Data & Code Availability (Not applicable)**

| Description | Source / Repository | Persistent ID / URL |
|---|---|---|

**Other**

| Description | Source / Repository | URL / Catalog Number |
|---|---|---|
| **Software** | | |
| GraphPad Prism (10.2.3) | GraphPad Software | https://www.graphpad.com/features |
| PyMOL (4.6.0) | Schrodinger, LLC | https://pymol.org/ |
| UCSF Chimera (1.17.3) | 2018 Regents of the University of California | https://www.cgl.ucsf.edu/chimera/ |
| Adobe Photoshop 2024 | Adobe | https://www.adobe.com/products/photoshop/ |
| RaptorX | University of Chicago | http://raptorx6.uchicago.edu/ |
| SnapGene | GSL Biotech LLC | https://www.snapgene.com/ |
| NIH ImageJ | Research Services Branch (RSB) of the National Institute of Mental Health (NIMH) | https://imagej.net/nih-image/ |
| AutoDock Vina (1.2.0) | Center for Computational Bioinformatics for the Scripps Research Institute | https://vina.scripps.edu/ |
| FlowJo v10.10 | BD Biosciences | https://www.flowjo.com/solutions/flowjo |

| **Drugs/Agents/Kits/Equipment Used** | | |
|---|---|---|
| **Description** | **Source** | **Catalog Number** |
| 0.2 uM syringe filter | VWR | 28145-477 |
| 150 mm Tissue Culture plate | VWR | 734-2754 |

| | | |
|---|---|---|
| 16% PFA | Electron Microscopy Science | 15710 |
| 1X HBSS | ThermoFisher Scientific | 14025092 |
| 1X PBS | Thermo Scientific | 14190144 |
| 2-β-mercaptoethanol | J.T. Baker | 4049-00 |
| 24-well cell culture plate | Cellstar | 662160 |
| 30% Acrylamide/Bis Solution, 37.5:1 | BioRad | 1610158 |
| 70 μM Cell Strainers for 1000 μL Pipette Tips | Fisher Scientific | 03-421-228 |
| Albumin test clips | IDEXX | 98-11065-01 |
| Allosteric PTP1B inhibitor | EMD Millipore | 539741-5MG |
| Ammonium Persulfate (APS) | Sigma | 215589 |
| Antibiotic-Antimycotic (100X) | Sigma | A5955 |
| Antibiotic-Antimycotic (100X) | ThermoFisher Scientific | 15240096 |
| BCA Protein Assay Kit | Thermo Scientific | 23227 |
| Bead beater | DENTSPLY | C321001 |
| Benzamine Hydrochloride | Sigma | 63226-1mL-F |
| Bio-Rad Low-Pressure Chromatography Pump EP-1 | BioRad | N/A |
| BioPulverizer | Biospec Products | 59012N |
| Biotinylated Insulin | Eagle Bioscience | INS30-G100 |
| BL21 (DE3) maximum efficiency *E. coli* competent cells | ThermoFisher Scientific | EC0114 |
| Blasticidin S HCl | Gibco | A11139-03 |
| Blotting-grade blocker (nonfat dry milk) | BioRad | 1706404 |
| BMG Labtech | FlourStar Omega | N/A |
| Bovine Serum Albumin | Roche | 10775835001 |
| Bovine thrombin | BioPharm Laboratories, LLC | PN: 91-010-005 |
| Bromphenol blue | Sigma | B0126 |
| BSA | Roche | 10775835001 |
| BUN test clips | IDEXX | 98-11070-01 |
| Carbenicillin | Sigma | C3416 |
| Cell culture incubator | Nuaire | NU-5810 |
| Chloroform (Molecular Biology Grade) | Fisher Scientific | AAJ67241AP |
| Collagen Solution | Sigma | C4243 |
| cOmplete, EDTA Free Protease inhibitor tablet | Roche | 11873580001 |
| Confocal Microscope | Olympus | FV100 |
| control siRNA | ThermoFisher Scientific | 4390844 |
| Corning® 12 mm Transwell® with 0.4 µm Pore Polyester Membrane Insert, Sterile | Corning | 3460 |
| Creatine test clips | IDEXX | 98-11074-01 |
| Cycloheximide (CHX) | Sigma | 01810 |
| DAPI | Sigma | D9542 |
| DEPC-treated nuclease-free water | Thermo Fisher | AM9906 |
| Detergent-compatible (DC) Protein Assay | BioRad | 5000111 |
| dithiothreitol | Sigma | 10197777001 |
| DMSO | Acros | 327182500 |
| Donkey Serum | JacksonImmuno | 017-000-001 |

| | | |
|---|---|---|
| DTT | Sigma | 10197777001 |
| Dulbecco's Modified Eagle Medium (DMEM) | Gibco | 11960-044 |
| EBM-2 | Lonza | CC-3156 |
| EDTA | Thermo Fisher | 15575020 |
| EDTA-free protease inhibitor cocktail | Sigma | 11836170001 |
| EGM-2 | Lonza | CC-4176 |
| EGTA | EMD Millipore | 324626 |
| Electron Microscopy Sciences Round Cover Slip German Glass #1.5, 8 mm | FisherScientific | 50-949-314 |
| Falcon® 100 mm TC-treated Cell Culture Dish, 20/Pack, 200/Case, Sterile | Corning | 353003 |
| Fetal bovine serum (FBS) | Fisher Scientific | 3SH30910.03 |
| Flourecent glucose analog 2-[N-(7-nitrobenz-2-oxa1,3-diazol-4-yl) mino]-2-deoxy-d-glucose (2-NBDG) | ThermoFisher Scientific | N13195 |
| Gibco™ Opti-MEM™ I Reduced Serum Medium | Fisher Scientific | 31-985-070 |
| glucose-free DMEM | ThermoFisher Scientific | 11966025 |
| Glutathione Sepharose 4B | Cytiva | 17075601 |
| Glycerol | Acros | 15892-0025 |
| heparin sodium salt | Sigma | H3393 |
| HEPES | Sigma | H3375 |
| HNMPA-(AM)3 | Santa Cruz Biotechnology | sc-221730 |
| iBlot transfer stacks | Thermo Scientific | IB23001 |
| iBright CL750 imaging system | Thermo Scientific | A44116 |
| Igepal CA-630 | Sigma | 56741 |
| Immulon 4 HBX ELISA Plates | Thermo Scientific | 3855 |
| Imperial Protein Stain | ThermoFisher Scientific | 24615 |
| Invitrogen™ iBlot™ 2 Gel Transfer Device | Fisher Scientific | IB21001 |
| iScript cDNA Synthesis Kit | BioRad | 1708890 |
| Isopropyl-B-D-thiogalactoside (IPTG) | Roche | 11422446001 |
| $K_2$EDTA Vials | Tiger Medical | TM28797 |
| KCl | Sigma | P3911 |
| Leupeptin | Sigma | L2884 |
| Lipofectamine 2000 | ThermoFisher Scientific | 11668019 |
| Lipofectamine RNAiMAX | Invitrogen | 56532 |
| LY294002 | CST | 9901S |
| Malachite green solution | Echelon Biosciences | K-1501 |
| $MgCl_2$ | Sigma | M8266 |
| Mouse antithrombin III | Haematologic Technologies | MCATIII-5120 |
| Mouse thrombin | Molecular Innovations | MTHROM |
| N-acetyl-L-leucyl-L-leucyl-L-norleucinal | Sigma | 11086090001 |
| NEB Stable competent *E. coli* (High Efficiency) | NEB | C3040H |
| Nuclease-free Tris pH 7.5 | BioWorld | 21420063-3 |
| *Par1* siRNA | ThermoFisher Scientific | 4390824-s4923 |
| Peroxidase Streptavidin | Jackson Immuno | 016-030-084 |

| | | |
|---|---|---|
| Phosphatase Substrate | Echelon Biosciences | 871-70 |
| Phosphate Standard | Upstate Cell Signaling Solutions | 20-103 |
| Plate Washer | BioRad | Model 1550 |
| PMSF | Sigma | P7626 |
| Polybrene | Sigma | TR-1003-G |
| Propidium iodide | BD Biosciences | 556463 |
| PTP1B enzyme | EMD Millipore | KP8401-5UG |
| PTP1B Inhibitor II (JTT-551) | EMD Millipore Corp | 5.30821.0001 |
| RNaseOUT | Thermo Fisher | 10777019 |
| RNeasy Mini Kit | Qiagen | 74106 |
| S961 Acetate | MedChemExpress | HY-P2093B |
| Saracatinib | Selleckchem | AZD0530 |
| SDS | Sigma | L5750 |
| Serum Vial [clot activator/SST™ Gel (Amber)] | Tiger Medical | 365978 |
| Sigma Delta Vaporizer | Penlon | 52606 |
| Sodium fluoride | Sigma | S6521 |
| Sodium molybdate | Sigma | M1003 |
| Sodium orthovanadate | MP Biomedical | 159664 |
| Sodium pyrophosphate | Sigma | 221363 |
| SsoAdvance Universal SYBR Green Supermix | BioRad | 1725275 |
| Streptavidin | NEB | N7021S |
| Sulfuric Acid (6N) Standardized Solution | Thermo Scientific | 035610.K2 |
| SuperSignal™ West Femto Maximum Sensitivity Substrate | Thermo Scientific | 37075 |
| T25 flask | ThermoFisher Scientific | 156367 |
| T75 Flask | Cellstar | 658-170 |
| Tamoxifen | Sigma | T5648 |
| TEMED | Santa Cruz | sc-29111 |
| Terrific Broth | ThermoFisher Scientific | 22711022 |
| TMB | Thermo Scientific | N301 |
| Tris-HCl | Roche | 10812846001 |
| Triton X-100 | ThermoFisher Scientific | A16046.AE |
| Trypsin | ThermoFisher Scientific | 25200056 |
| Tween 20 | VWR | 0777-1L |
| Tween-20 | VWR | 0777-1L |
| UltraClear microscope slides | Denville Scientific Inc | 14634019 |
| Vorapaxar | MedChemExpress | HY-10119/C |
| West Dura ECL | Thermo Scientific | 34076 |
| West Femto ECL | Thermo Scientific | 34095 |
| ZOE Fluorescent Cell Imager | BioRad | 1450031 |

### SUPPLEMENTARY APPENDIX (ONLINE FILES)

Supplemental File 1: (SF1.DNA_Sequences_FASTA) DNA sequences of all constructs used in the study in FASTA formatting.

Supplemental File 2: (SF2.Data) Excel file of all data values generated in the study. Each figure subpanel is listed as a separate tab.

Supplemental File 3: (SF3.Stats-Gender_Tables) Excel file of all statistical tests used in the study and list of all mice and genders.

Supplemental File 4: (SF4.Raw_Blots) Copy of all unedited blots displayed in the manuscript

### ARRIVE GUIDELINES

The ARRIVE guidelines (https://arriveguidelines.org/) are a checklist of recommendations to improve the reporting of research involving animals. Key elements of the study design should be included below to better enable readers to scrutinize the research adequately, evaluate its methodological rigor, and reproduce the methods or findings.

**Study Design:**

| **Groups** | **Sex** | **Age** | **Number (prior to experiment)** | **Number (after termination)** | **Littermates (Yes/No)** |
|---|---|---|---|---|---|
| Control (Cre-negative) | M | 3-6 months | Not available | 54 | Yes |
| $Par1^{iECko}$ | M | 3-6 months | Not available | 9 | Yes |
| $Par4^{iECko}$ | M | 3-6 months | Not available | 5 | Yes |
| $Par1/4^{iECko}$ | M | 3-6 months | Not available | 42 | Yes |
| $Par1^{iEChet};Par4^{iECko}$ | M | 3-6 months | Not available | 2 | Yes |
| $Par1/4^{iECko};Insr^{iEChet}$ | M | 3-6 months | Not available | 17 | Yes |
| $Par1/4^{iECko}$ - RiboTag | M | 3-6 months | Not available | 4 | Yes |
| $Par1^{iEChet};Par4^{iECko}$- RiboTag | M | 3-6 months | Not available | 6 | Yes |
| RiboTag | M | 3-6 months | Not available | 9 | Yes |

**Sample Size: Please explain how the sample size was decided. Please provide details of any a *priori* sample size calculation, if done.**

No specific criteria were used to determine the sample size for each experiment.

**Inclusion**

All mice with appropriate genotypes and ages were included in the experiments.

**Exclusion Criteria**

No mice were excluded from the study.

**Randomization**

Mutant mice were randomly assigned to experimental groups.

**Blinding**

All mice were assigned an alphanumeric value, and the operator was blinded to genotype while analyses were performed. Data were collated by genotype only after analyses were performed.
